# The UPR^ER^ governs the cell-specific response of human dopaminergic neurons to mitochondrial stress

**DOI:** 10.1101/2024.06.17.599325

**Authors:** Jana Heneine, Claire Colace-Sauty, Christiane Zhu, Benjamin Galet, Justine Guégan, François-Xavier Lejeune, Thomas Gareau, Noemi Asfogo, Corinne Pardanaud-Glavieux, Olga Corti, Philippe Ravassard, Hélène Cheval

**Affiliations:** Institut du Cerveau - Paris Brain Institute - ICM, Sorbonne Université, Inserm U1127, CNRS UMR7225, APHP, Hôpital Pitié Salpêtrière, Paris, France

**Keywords:** human dopaminergic neurons, UPR^ER^, PERK, lncRNAs, mitophagy, mitochondrial DNA biogenesis, translation

## Abstract

Mitochondrial dysfunction is thought to be central to the pathophysiology of Parkinson’s disease. The preferential vulnerability of dopaminergic (DA) neurons of the *substantia nigra pars compacta* to mitochondrial stress may underlie their massive degeneration and the occurrence of motor symptoms. Using LUHMES-derived DA neurons, we demonstrated that inhibition of the mitochondrial electron transport chain resulted in a severe alteration of mitochondrial turnover, pushing the balance towards mitochondrial loss, a reduction of the maturation status of the DA population and an increased proportion of apoptotic cells. PERK-mediated Unfolded Protein Response of the Endoplasmic Reticulum (UPR^ER^) emerged as the key coordinator of the stress response, governing the inactivation of the mitochondrial UPR (UPR^mt^), the initiation of mitophagy and the cell-specific expression of long non-coding RNAs (lncRNAs). Importantly, we discovered novel lncRNAs specifically expressed in human DA neurons upon stress. Among them, we showed that lnc-SLC6A15-5 contributes to the resumption of translation after mitochondrial stress.

**Summary:** The Unfolded Protein Response of the Endoplasmic Reticulum is induced upon stress in human dopaminergic neurons and modulates mitochondrial homeostasis and transcriptional programs including expression of long non-coding RNAs (lncRNAs). We discovered a lncRNA involved in translation resumption after stress.

## Introduction

Parkinson’s Disease (PD) is a prevalent neurological disorder characterized by the degeneration of several neuronal subtypes, but affecting predominantly the dopaminergic (DA) neurons of the *substantia nigra pars compacta* (SNpc) (Pacelli *et al*, 2015; Brichta & Grelnengard, 2014; Pissadaki & Bolam, 2013; Damier *et al*, 1999; Hirsch *et al*, 1988). The progressive and massive DA neuronal loss constitutes a hallmark of the disease, responsible for the major motor symptoms observed in patients, rigidity, bradykinesia and tremor (Dickson *et al*, 2009; Kalia & Lang, 2015). Mitochondrial dysfunction has emerged as a prominent player in PD pathogenesis, marked by several lines of evidence in PD patients and animal models. Defects in mitochondrial complex I activity and mitochondrial DNA homeostasis have been shown in brain tissue from PD patients (Schapira *et al*, 1989; Dölle *et al*, 2016; Borsche *et al*, 2021; Grünewald *et al*, 2019), and exposure to environmental mitochondrial toxins, such as 1-methyl-4-phenyl-1,2,3,6-tetrahydropyridine (MPTP) or rotenone, has been linked to the manifestation of clinical symptoms resembling PD and underlying DA neurodegeneration in the SNpc (Langston *et al*, 1983; Sherer *et al*, 2003a, 2003b). Furthermore, an ever growing number of studies reported changes to mitochondrial biology using various cellular and animal models (Dauer & Przedborski, 2003; Pacelli *et al*, 2015; Bose & Beal, 2016). Importantly, the significance of mitochondrial alterations in the pathophysiology of PD has been emphasized by the discovery of the causal link between mutations in *PRKN* and *PINK1* and autosomal recessive forms of PD (Pickrell & Youle, 2015). *PRKN* and *PINK1* encode the E3 ubiquitin ligase PARKIN and the mitochondrial serine/threonine kinase PINK1, which hold joint pivotal roles in mitochondrial quality control in response to mitochondrial dysfunction (Eldeeb *et al*, 2022; Zhu *et al*, 2013). Altogether, functional changes to mitochondria naturally accumulating during aging are suspected to lead to a homeostatic imbalance that significantly enhances the vulnerability of DA neurons of the SNpc to cell death compared to other neuronal subtypes. The selective effects of *PRKN* and *PINK1* gene mutations, which sensitize primarily the SNpc DA neurons to cell death, despite being ubiquitously expressed in various cell types, raises intriguing questions about the DA neuron-specific factors contributing to mitochondrial stress vulnerability.

So far, specific molecular signatures defining neuronal cells have been obtained using transcriptomic data focused on protein-coding genes. However, non-coding elements of the genome, such as long non-coding RNAs (lncRNAs), are gaining prominence for their cell-specific regulatory functions, spanning from epigenetic to post-translational levels (Cabili *et al*, 2011; Morán *et al*, 2012; Washietl *et al*, 2014; Ward *et al*, 2015; Jiang *et al*, 2016; Akerman *et al*, 2017; Liu *et al*, 2017; Gendron *et al*, 2019; Seifuddin *et al*, 2020; de Goede *et al*, 2021; Ulitsky & Bartel, 2013; Jarroux *et al*, 2017; Mattick *et al*, 2023). Moreover, most lncRNAs exhibit limited conservation across species and the vast majority of PD-associated single nucleotide polymorphisms (SNPs) fall into non-coding regions with potential regulatory functions (Altshuler *et al*, 2008; Nalls *et al*, 2019). Consequently, lncRNAs, along with their associated molecular mechanisms, emerge as promising candidates for elucidating the specific molecular mechanisms underlying vulnerability to stress and, by extension, the pathophysiology of human diseases associated with the alteration of specific cellular subtypes. Due to their weak inter-species conservation, their relevance takes on an even more significant dimension in the context of pathologies for which animal models do not fully recapitulate the human clinical manifestations, such as PD.

In the context of mitochondrial stress, extensive research has unveiled the central role of the integrated stress response (ISR) in human cells (Krug *et al*, 2014; Quirós *et al*, 2017; Jennings *et al*, 2023; van der Stel *et al*, 2022; Carta *et al*, 2023), leading to the activation of the PERK-mediated Unfolded Protein Response of the Endoplasmic Reticulum (UPR^ER^). PERK-dependent phosphorylation of EIF2α results in the attenuation of general translation, while allowing for selective translation of stress-associated proteins, such as ATF4, which initiates key transcriptional programs promoting pro-survival or pro-apoptotic responses, depending on the severity and duration of the stress (Wek & Cavener, 2007). However, the extent to which other branches of the UPR^ER^, mediated by the activation of IRE1 or ATF6, or the UPR^mt^, are involved in coping with mitochondrial stress is still unclear and highly differ across different cell types and mitochondrial stress conditions (Quirós *et al*, 2017; Cai *et al*, 2020).

In this study, we demonstrate that exposing human DA neurons derived from LUHMES cells (Lotharius *et al*, 2002; Scholz *et al*, 2011) to inhibitors of the electron transport chain prompted the simultaneous activation of all branches of the UPR^ER^, with a pronounced emphasis on the PERK-UPR^ER^ pathway. This latter pathway contributed to induction of stress-induced mitophagy and inactivation of the UPR^mt^ in neurons. Importantly, we discovered novel lncRNAs expressed in DA neurons specifically during the mitochondrial stress response, downstream of the PERK-mediated UPR^ER^. Among these stress-induced lncRNAs, *lnc-SLC6A15*, emerged as a regulator of translation resumption that occurs following mitochondrial stress.

## Results

### Mitochondrial stress induced by the inhibition of the electron transport chain destabilizes mitochondrial turnover in human DA neurons

To study the effect of mitochondrial stress on human DA neurons, we used DA neurons generated from LUHMES cells (LUnd Human MESencephalic neuronal cell line, immortalized DA progenitors; Lotharius *et al*, 2002; Scholz *et al*, 2011). LUHMES cells differentiate rapidly and homogeneously into DA neurons that can be produced in large numbers, facilitating PD research (Lotharius *et al*, 2002, 2005; Höllerhage *et al*, 2017; Pierce *et al*, 2018). Accordingly, after 6 days of differentiation, 90% of the cells expressed the enzyme tyrosine hydroxylase (TH), essential to the production of dopamine, and 74% co-expressed TH with the DA transporter DAT **(Figure 1a**; **Figure 2a, b, d)**. At this time point, we treated the neurons for 8 h with the mitochondrial toxins antimycin A and oligomycin, which trigger mitochondrial stress through the inhibition of the complex III and the ATP synthase of the mitochondrial respiratory chain respectively. Importantly, these toxins have been shown to induce PINK1/PARKIN-dependent mitophagy, a mitochondrial quality control mechanism relevant to PD (Lazarou *et al*, 2015; Georgakopoulos *et al*, 2017).To confirm its activation, we assessed the phosphorylation of ubiquitin at Serine 65, a marker of the early phase of this mitophagy program (Kazlauskaite *et al*, 2014; Wauer *et al*, 2015; Ge *et al*, 2020; Picca *et al*, 2021). We observed staining for phosphorylated ubiquitin in around 60% of the neurons as early as 4 h after application of these toxins, compared to 2 to 5% in control conditions (**Figure 1b, c)**. Mitophagy initiation was associated with an alteration of the mitochondrial network reminiscent of mitochondrial fragmentation, as demonstrated by the scattered localization of the mitochondrial import receptor subunit TOMM20 upon stress compared to the control neurons exhibiting a rather clustered TOMM20 staining (**Figure 1b, d**).

**Figure 1.**
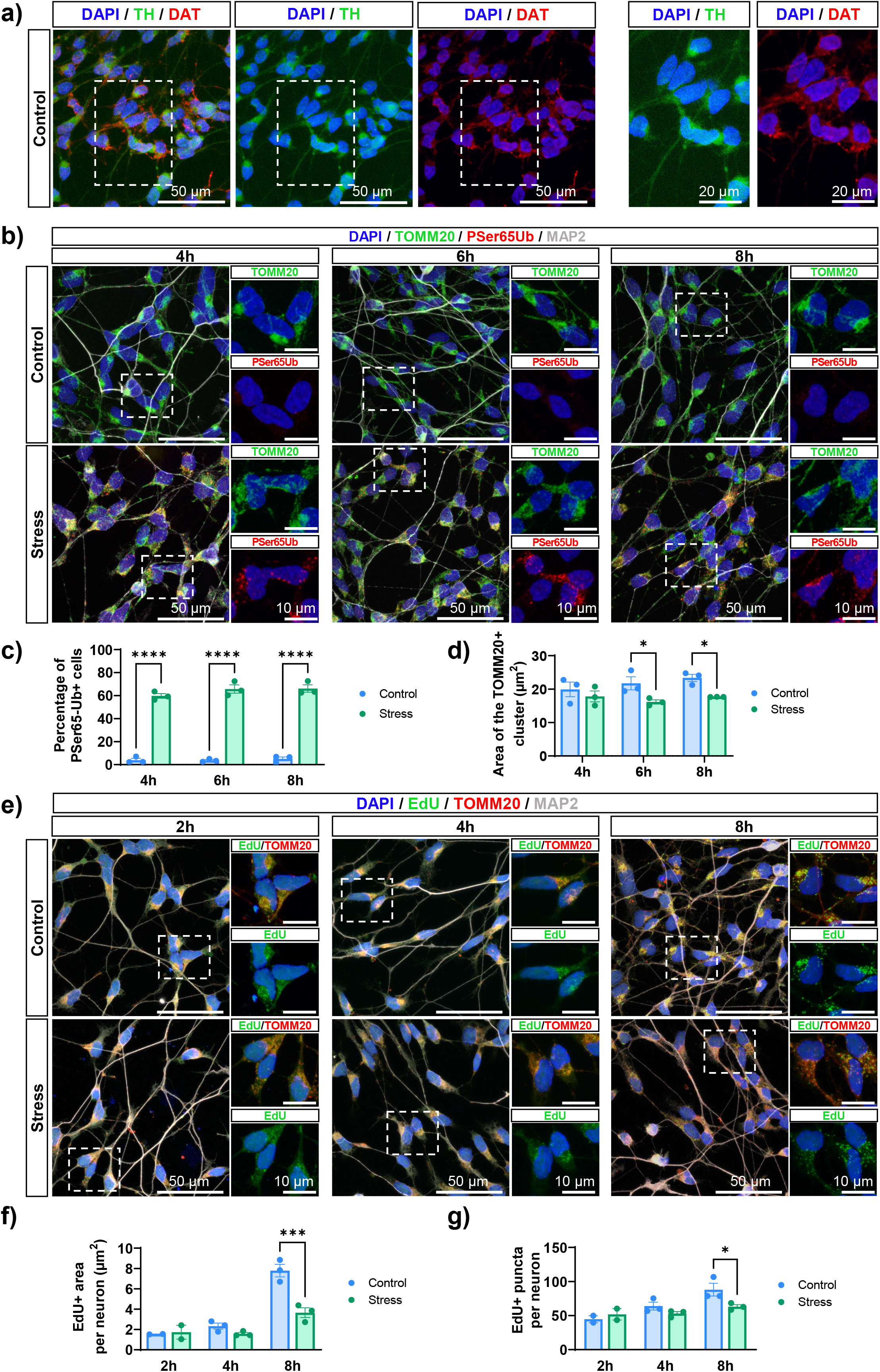
Inhibition of the mitochondrial electron transport chain induces mitophagy and a decrease in mitochondrial biogenesis in DA neurons. **(a)** TH (green) and DAT (red) expression, assessed by immunofluorescence on LUHMES cells differentiated for 6 days. **(b)** Phospho-Serine 65 ubiquitin (red), TOMM20 (green) and MAP2 (grey) expression in DA neurons treated with DMSO (Control) or after exposition to mitochondrial toxins for 4 h, 6 h and 8 h (Stress), observed by immunofluorescence. **(c)** Percentage of phospho-Serine65 ubiquitin-positive neurons in control conditions and following 4 h, 6 h or 8 h of mitochondrial stress (Two-way ANOVA with Tukey’s multiple comparisons test). **(d)** Area (in µm^2^) of the TOMM20-positive cluster in neurons in control conditions and following 4 h, 6 h or 8 h of mitochondrial stress (Mixed-effects analysis withTukey’s multiple comparisons test). **(e)** MAP2 (grey) and TOMM20 (red) expression assessed by immunofluorescence and EdU (green) detection in control conditions or following mitochondrial stress for 2 h, 4 h or 8 h. **(a, b and e)** Nuclei were stained using DAPI (blue). For each low magnification photograph, areas indicated by dotted lines are zoomed and presented on the right panel. **(f)** Area comprising EdU signal (in µm^2^) per neuron in control conditions or following mitochondrial stress for 2 h, 4 h or 8 h (Mixed-effects analysis and Bonferroni’s multiple comparisons test). **(g)** Number of EdU-positive puncta per neuron in control conditions and upon a 2 h-, 4 h- or 8 h-stress (Mixed-effects analysis and Bonferroni’s multiple comparisons test). **(c, d, f and g)** Each dot represents the data obtained from 3 independent differentiation experiments. The bar represents the mean of the 3 values, and the error bars show standard error of the mean. *p-value ≤ 0,05, **** p-value ≤ 0,0001

**Figure 2.**
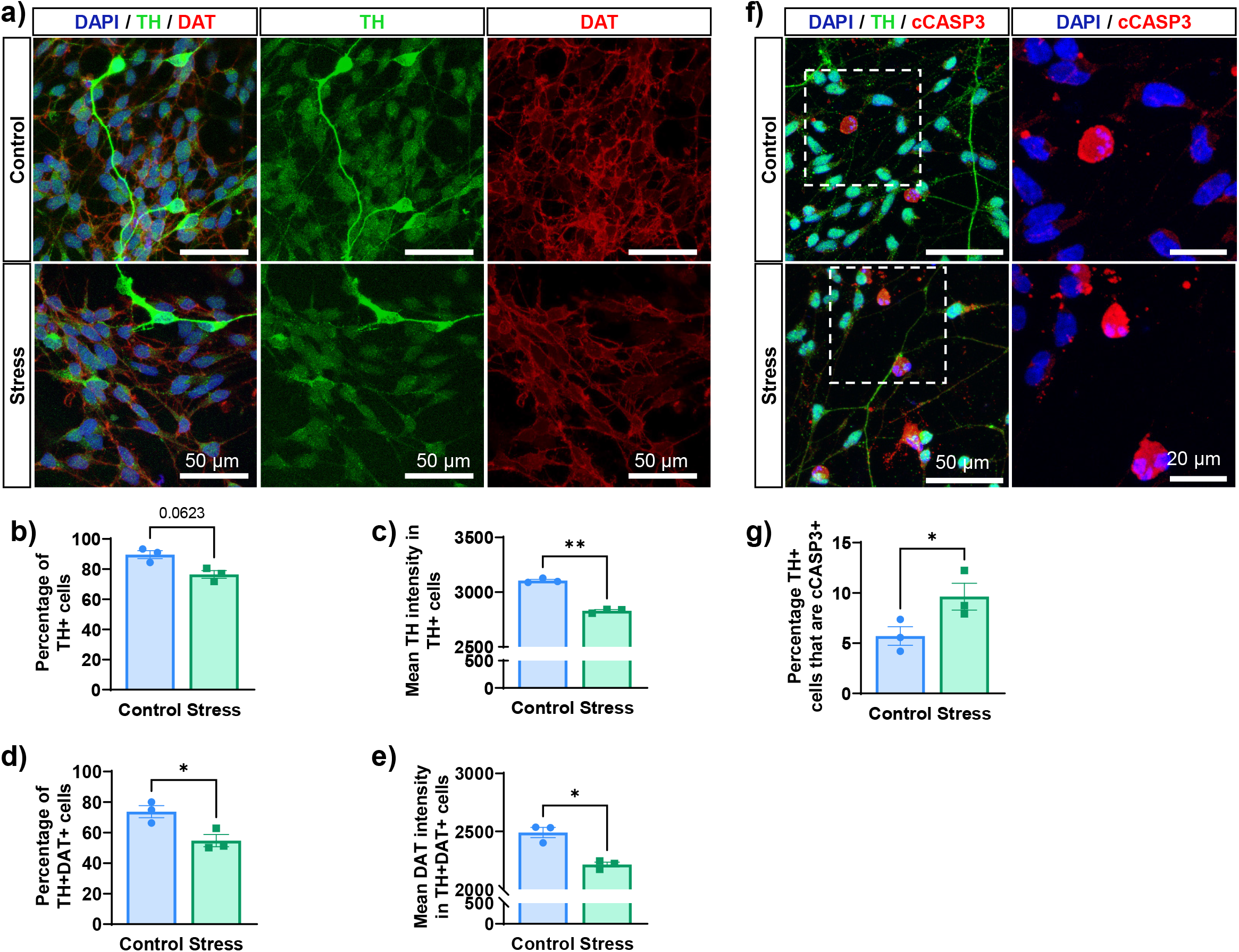
Mitochondrial stress leads to loss of mature dopaminergic marker and cell death. **(a)** TH (green) and DAT (red) expression, assessed by immunofluorescence on DA neurons in control conditions or upon 8 h mitochondrial stress. Quantification of the **(b)** percentage of TH^+^ cells (two-tailed paired t-test) **(c)** mean TH signal intensity per TH^+^ cell (two-tailed paired t-test), **(d)** percentage of TH^+^ DAT^+^ cells (two-tailed paired t-test) and **(e)** mean DAT signal intensity per TH^+^ DAT^+^ cell (two-tailed paired t-test) in both control and stress (8 h) conditions. **(f)** TH (green) and cleaved CASP3 (cCASP3, red) expression, assessed by immunofluorescence on DA neurons in control conditions or upon 8 h-mitochondrial stress. **(a and f)** Nuclei were stained using DAPI (blue). For each low magnification photograph, areas indicated by dotted lines are zoomed and presented on the right panel. **(g)** Quantification of the percentage of TH^+^ cells that exhibited cCASP3^+^ staining in both control and stress conditions (two-tailed paired t-test). **(b, c, d, e and g)** Each dot represents the data obtained from 3 independent differentiation experiments. The bar represents the mean of the 3 values, and the error bars show standard error of the mean. *p-value ≤ 0,05, **p-value ≤ 0,01.

Mitochondrial turnover in physiological conditions as well as under stress relies on the fine balance between mitophagy and mitochondrial biogenesis (Zhu et al., 2013). We therefore investigated *de novo* synthesis of mitochondrial DNA (mtDNA) as a marker of mitochondrial biogenesis by studying the incorporation of the thymidine analog EdU. As mature neurons are post-mitotic cells, EdU integration is expected to be specific to mtDNA (Prole *et al*, 2020). Consistently, we observed co-localization of 90% of the EdU puncta with TOMM20 (**Supplementary Figure 1b-c**) and massive reduction of the EdU signal after treatment of the neurons with 2’,3’-dideoxycytidine (ddC), an inhibitor of chain elongation (**Supplementary Figure 1a**). Following 8 h of treatment with mitochondrial toxins, the area occupied by the EdU signal was reduced by half compared to control conditions, and the number of EdU-positive puncta per neuron was decreased by 30%, with no change at earlier time points (**Figure 1e-g**), demonstrating stalled synthesis of mtDNA. Altogether, these results indicate impairment of mitochondrial turnover following stress in DA neurons, with an overall induction of mitophagy and decrease of mitochondrial biogenesis.

### Mitochondrial stress alters the maturity and survival rate of human DA neurons

We then examined whether the inhibition of the electron transport chain also affected the identity and survival of LUHMES-derived DA neurons. We observed a tendency towards a reduction in the percentage of DA neurons expressing TH upon stress compared to control conditions (90 % in controls *versus* 77% upon stress; p= 0,0623; **Figure 2a, b**), with an overall decrease in TH signal intensity (**Figure 2c**). Treatment with the mitochondrial toxins also led to a 26% decrease in the percentage of mature DA neurons expressing both TH and DAT (**Figure 2d**) and in a reduction of DAT signal intensity in DA neurons with residual expression (**Figure 2e**). These results suggest that mitochondrial stress induced an alteration of the DA neurons maturation status. In parallel, we observed that the proportion of DA neurons expressing the pro-apoptotic marker cleaved Caspase 3 (cCASP3) slightly rose from 5% in control conditions to 10% following mitochondrial stress, indicating increase in cell death (**Figure 2f, g**).

### Mitochondrial stress leads to the concomitant activation of the three UPR^ER^ branches and inhibition of neuronal development pathways in human DA neurons

To decipher the signaling pathways induced in DA neurons in response to mitochondrial stress, we investigated the stress-associated alterations of the transcriptome using RNA-seq. Principal component analysis demonstrated that datasets generated from DA neurons treated with mitochondrial toxins or with vehicle (DMSO) alone formed two distinct clusters (**Figure 3a**). Application of mitochondrial stress accounted for nearly 60% of the variance between the samples, as shown at the PC1 level, represented on the x axis. Focusing on protein-coding genes, we identified 12898 unique transcripts, including 772 genes with significant upregulation of expression upon stress and 605 with significant downregulation. Gene ontology analysis on the latter category revealed an enrichment in genes associated with the biological process “Nervous system development” (**Figure 3b**), reminiscent of the significant decrease in mature DA neurons observed following mitochondrial stress (**Figure 2a-e**). In addition, mRNA levels of all the 13 mitochondrial genes encoding sub-units of complexes I, III, IV and V of the electron transport chain (Schon *et al*, 2012), were significantly reduced (**Figure 3c**). Together with the alterations in mitochondrial DNA synthesis (**Figure 1e-g**), this indicated that mitochondrial genomic programs are strongly impaired in response to the treatment with mitochondrial toxins. In parallel, gene ontology analysis on the 772 protein-genes upregulated upon stress revealed a strong enrichment in genes associated with the Unfolded Protein Response of the endoplasmic reticulum (UPR^ER^) with the terms “Response to Endoplasmic Reticulum Stress”, “Intrinsic Apoptotic Signaling Pathway in Response to ER Stress” and “Response to Unfolded Protein” (**Figure 3b**). Interestingly, many terms from this analysis also referred to amino acid transport processes, previously shown to be induced via the integrated stress response, in particular the UPR^ER^-PERK pathway, during ER stress (Harding *et al*, 2003; Han *et al*, 2013; Quirós *et al*, 2017). Several studies have highlighted the role of the UPR^ER^ in response to mitochondrial stress in mammalian cells (Quirós *et al*, 2017; Krug *et al*, 2014; Jennings *et al*, 2023; van der Stel *et al*, 2022; Carta *et al*, 2023), and show a predominant role of the PERK-ATF4 pathway, with no or weak activation of the other UPR^ER^-associated branches, i.e. IRE1-XBP1 or ATF6 pathways. In contrast, pathway analysis on our datasets revealed concomitant activation of all branches of the UPR^ER^ at the transcriptional level upon stress (**Figure 3d**). To validate this result and assess the activation kinetics of the different UPR^ER^ pathways, we examined mRNA expression of several key players in each pathway at different time points during the treatment with the mitochondrial toxins compared to the control condition (**Figure 3e**). The PERK-EIF2α-mediated UPR^ER^ was strongly activated as early as 2 h into toxin exposure, as shown by the overexpression of *ATF4* mRNA and its target genes, i.e. *ATF3*, *DDIT3*, *TRIB3* and *CHAC1*, which are involved in cell death programs (Han et al., 2013). In parallel, the early overexpression of *NRF2* mRNA indicates that PERK activation upon stress also resulted in induction of the signaling pathway dependent on the antioxidant factor NRF2. We confirmed the activation of the IRE1-mediated UPR^ER^ upon stress, as demonstrated by the increased expression of *XBP1s*, generated by IRE1-dependent splicing of *XBP1,* as early as 2 h of stress (Park *et al*, 2021), and by the overexpression of XBP1s target gene, *DNAJC3* after 6 h of stress. In contrast, the IRE1-dependent TRAF2-JUNK pathway was not induced. Regarding the ATF6 pathway, we showed an upregulation of its target genes *HSPA5* and *XBP1*, after 30 min and 4 h of stress respectively. Moreover, expression of *EDEM1* and *HERPUD1*, encoding proteins involved in ER-associated degradation (ERAD) and associated with both IRE1- and ATF6-mediated UPR^ER^ pathways (Adachi *et al*, 2008; Park *et al*, 2021), was increased from 2 to 4 h of treatment with the mitochondrial toxins. Of note, PERK, IRE1 and ATF6 activities are known to be regulated by post-translational modifications upon stress, allowing for the induction of a rapid response and explaining the absence of early transcriptional changes for these genes upon stress. However, we observed that *ATF6* and *PERK* mRNA expression was upregulated at 4 h and 8 h of stress respectively, suggesting adaptations in the stress response across time. As expected with the activation of PERK (Han *et al*, 2013), there was a 2,6 fold increase in the phosphorylation of EIF2α upon stress (**Figure 3f**), indicating attenuation of general translation. Interestingly, none of the branches of the mitochondrial UPR (UPR^mt^) appeared to be involved in the stress response at the investigated time points (**Figure 3e**), as previously shown (Quirós *et al*, 2017). Except for the up-regulation of the ATF4-target gene *ATF5*, expression of the associated chaperones *YME1L1*, *LONP1* and *CLPP* was unchanged or even decreased upon stress (**Figure 3e**, lower panel). Similarly, there was no change or a tendency towards reduced expression for key genes of the SIRT3 UPR^mt^ pathway (i.e. *FOXO3* and *SIRT3*) and *NRF1*, a target gene of the ERα-mediated UPR^mt^. As a whole, we established that mitochondrial stress in human DA neurons triggered transcriptional programs responsible for loss of neuronal identity and activation of the PERK-, IRE1 and ATF6-mediated UPR^ER^, leading to an engagement towards apoptosis.

**Figure 3.**
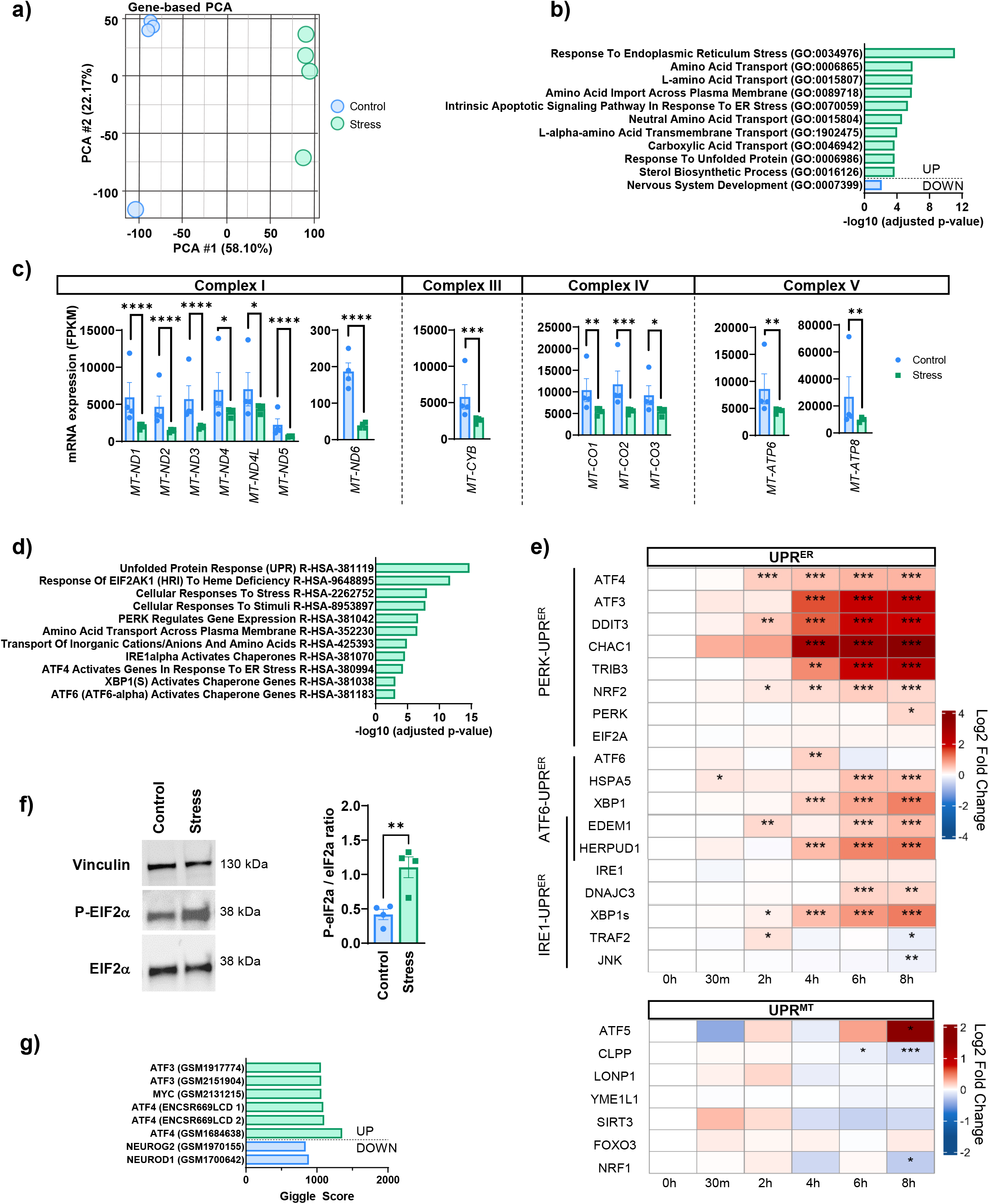
Mitochondrial stress triggers the Unfolded Protein Response of the Endoplasmic Reticulum (UPR^ER^) and inactivates neuronal development pathways. RNAseq datasets analysis were performed from 4 independent LUHMES cell differentiation experiments in control (DMSO) or stress conditions (oligomycin and antimycin) **(a)** Principal Component Analysis (PCA) of RNA-seq datasets from DA neurons in control (blue) or stress conditions (green). **(b)** Gene ontology analysis (Biological Process 2023, Enrichr) performed on protein-coding genes with upregulated (green) or downregulated (blue) expression after 8 h of mitochondrial stress compared to control conditions. **(c)** mRNA expression in FPKM of mitochondrial protein-coding genes in control or stress conditions using the RNA-seq datasets (Control, blue; Stress, green). The bar represents the mean of the 4 values per condition (Control, Stress), and the error bars show standard error of the mean (DESeq2 differential analysis). **(d)** Pathway analysis (Reactome 2022, Enrichr) performed on protein-coding genes with upregulated expression after 8 h of mitochondrial stress compared to control conditions. **(e)** Heatmaps representing mRNA expression by RT-qPCR of genes encoding main actors of the three branches of the UPR^ER^ and of the UPR^mt^ at different time points (30 min, 2 h, 4 h, 6 h and 8 h) during mitochondrial stress, compared to control conditions (Two-way ANOVA with Tukey multiple comparisons test). mRNA expression was normalized relatively to *TBP* mRNA expression. Data are represented in Log_2_(Fold change Stress/Control) for 3 independent experiments. **(f)** EIF2α, phosphorylated EIF2α (P-EIF2α) and vinculin expression assessed by western blot in control conditions and upon mitochondrial stress (8 h). Quantification of the P-EIF2α/EIF2α ratio in control conditions and upon mitochondrial stress (8 h) from 3 independent differentiation experiments represented by 3 dots (two-tailed paired t-test). The bar represents the mean of the 3 values, and the error bars show standard error of the mean. **(g)** Transcription factors with binding regions (established by publicly available ChIPseq datasets) showing a significant overlap with open chromatin regions associated with altered accessibility upon stress and determined by ATACseq on DA neurons (Cistrome DB). *p-value ≤ 0,05; **p-value ≤ 0,01; ***p-value ≤ 0,001; **** p-value ≤ 0,0001.

### The transcriptional response of human DA neurons to mitochondrial stress appears to rely primarily on the PERK-mediated UPR^ER^

We investigated further the regulation processes involved in the response of DA neurons to mitochondrial stress by studying changes in chromatin accessibility using ATAC-seq. This technology allows for the detection of potential active regulatory regions, such as promoters, enhancers, repressors etc. We identified 39720 peaks, reflecting regions of open chromatin, present in the 3 datasets obtained from the control cultures of DA neurons, and 39 375 peaks in the 4 datasets from DA neurons subjected to mitochondrial stress. We found 1327 regions more accessible and 2667 regions less accessible upon stress compared to control conditions (**Supplementary Figure 2a**). Most changes were observed in intragenic and intergenic regions, accounting for 85% and 63% of the regions respectively, with increased or in most cases decreased accessibility. In contrast to these latter categories, the number of promoter-associated regions with increased accessibility rose following mitochondrial stress. In line with the transcriptomic analyses, gene ontology enrichment analyses performed on the genes associated with these promoter regions confirmed the activation of transcriptional programs upon mitochondrial stress, in particular the engagement in the apoptotic pathway downstream of ER stress (**Supplementary Figure 2b**) and the alteration of neuronal identity (**Supplementary Figure 2c**).

Similarly, extending this analysis to all the regions with increased accessibility upon stress, we found enrichment in regions associated with gene regulation, stress response and apoptosis, whereas less accessible regions following stress were associated with cell-cell adhesion processes and nervous system development (**Supplementary Figure 2d**). To pinpoint key transcription factors involved in these genomic stress responses, we cross-referenced our data with ChiP-seq datasets and determined whether the identified stress-associated chromatin regions had already been experimentally shown to bind specific transcription factors (**Figure 2g**). Thus, regions found to be more accessible upon mitochondrial stress were significantly associated with ATF4, ATF3 as well as MYC, whereas regions with reduced accessibility were associated with transcription factors involved in neurodevelopment, such as NEUROD1 and NEUROG2.

Altogether, analysis of transcription factors binding sites within open chromatin regions suggested a predominant role for the PERK-EIF2a-ATF4 pathway in the mitochondrial stress response of DA neurons.

### PERK-mediated UPR^ER^ contributes to the regulation of mitochondrial turnover and the inhibition of the UPR^mt^ in human DA neurons exposed to mitochondrial stress

Given the highlighted prevailing contribution of the PERK-ATF4 UPR^ER^ pathway in the stress response of LUHMES-derived DA neurons and the close connection between the ER and mitochondria (Senft & Ronai, 2015), we assessed the direct involvement of this pathway in the observed mitochondrial alterations (**Figure 2**). To this end, we used the selective synthetic inhibitor GSK2606414 (compound 7-methyl-5-(1-{[3-(trifluoromethyl)phenyl]acetyl}-2,3-dihydro-1*H*-indol-5-yl)-7*H*-pyrrolo[2,3-d]pyrimidin-4-amine) to inhibit PERK (Axten *et al*, 2012; Mercado *et al*, 2018; Gundu *et al*, 2022) and examined the effects of this inhibition on mitophagy, mitochondrial biogenesis and the regulation of the UPR^mt^ upon stress. We first verified the expected effect of GSK2606414 on ATF4 expression following stress. As anticipated, in cells exposed to the mitochondrial toxins, there was a significant increase in ATF4 protein levels from 4 h to 8 h of treatment; this effect was similar to that of tunicamycin, an N-glycosylation inhibitor known to activate ER stress and the UPR^ER^ (**Figure 4a, b**). Addition of GSK2606414 to the medium significantly attenuated this response, reducing ATF4 protein levels in stressed cells to levels comparable to those in control cells. In addition, GSK2606414 inhibited the transcriptional induction of the ATF4 target gene *ATF3* following mitochondrial stress, confirming its overall inhibitory effect on the PERK-ATF4 pathway (**Figure 4c**).

**Figure 4.**
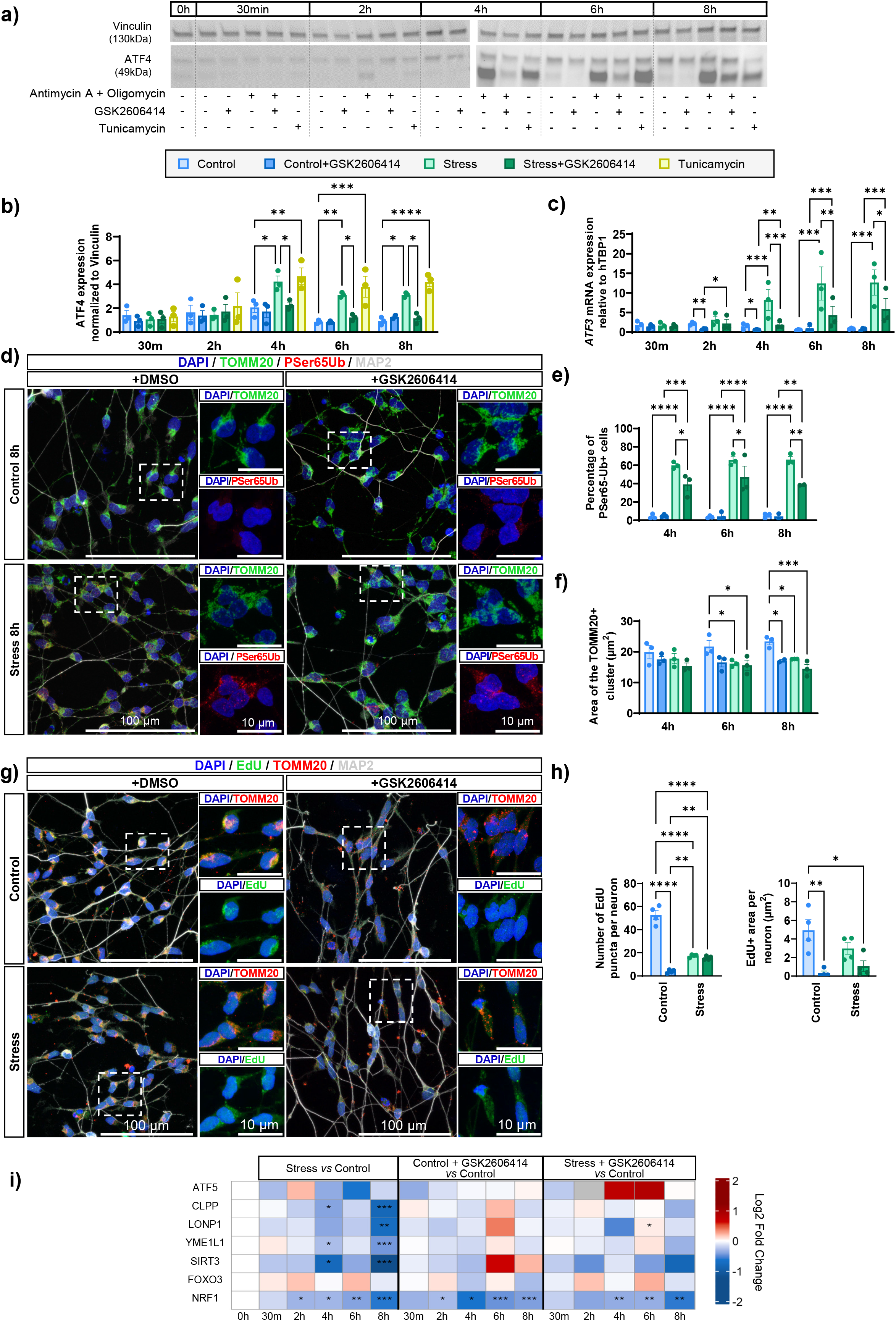
PERK-mediated UPR^ER^ regulates mitophagy and the UPR^mt^ upon stress, as well as the mitochondrial integrity and biogenesis at a basal level. **(a)** ATF4 and vinculin expression assessed by western blot in control conditions (DMSO only) and upon mitochondrial stress (antimycin A and oligomycin), in the presence or absence of GSK2606414, at different time points of stress (0 h, 30 min, 2 h, 4 h, 6 h and 8 h). Tunicamycin was used as a positive control for the activation of the PERK pathway. **(b)** Quantification of ATF4 protein expression (normalized to vinculin) in control conditions and upon mitochondrial stress, in the presence or absence of GSK2606414, at different time points of stress (0 h, 30 min, 2 h, 4 h, 6 h and 8 h). Data were normalized to the condition Control 0 h that was present in each gel (Two-way ANOVA with Tukey multiple comparisons test). **(c)** Quantification of *ATF3* mRNA expression assessed by RT-qPCR in control conditions and upon mitochondrial stress, in the presence or absence of GSK2606414, at different time points of stress (0 h, 30 min, 2 h, 4 h, 6 h and 8 h). mRNA expression was normalized relatively to *TBP* mRNA expression. **(d)** Phospho-Serine 65 ubiquitin (red), TOMM20 (green) and MAP2 (grey) expression in DA neurons treated with DMSO (Control) or after exposure to mitochondrial toxins for 8 h (Stress), in the presence or absence of GSK2606414, observed by immunofluorescence. **(e)** Percentage of phospho-Serine65 ubiquitin-positive neurons in control conditions and following 4 h, 6 h or 8 h of mitochondrial stress, in the presence or absence of GSK2606414 (Two-way ANOVA with Tukey multiple comparisons test). Data obtained in absence of GSK2606414 are shown in Figure 1b,c. **(f)** Area (in µm^2^) of the TOMM20-positive cluster in neurons measured in control conditions and after 4 h, 6 h or 8 h of mitochondrial stress, in the presence or absence of GSK2606414 (Mixed-effects analysis with Tukey multiple comparisons test). Data obtained in absence of GSK2606414 are shown in Figure 1b,d. **(b, c, e and f)** Data from 3 independent differentiation experiments, represented by 3 dots, were used. The bar represents the mean of the 3 values, and the error bars show standard error of the mean. **(g)** MAP2 (grey) and TOMM20 (red) expression assessed by immunofluorescence and EdU (green) detection in control conditions or following mitochondrial stress for 8 h, in the presence or absence of GSK2606414 (Two-way ANOVA with Tukey multiple comparisons test). **(d and g)** Nuclei were stained using DAPI (blue). For each low magnification photograph, areas indicated by dotted lines are zoomed and presented on the right panel. **(h)** Area comprising EdU signal (in µm^2^) per neuron and number of EdU-positive puncta per neuron in control conditions or following mitochondrial stress for 8 h (Two-way ANOVA with Tukey’s multiple comparisons test). Data were obtained from 4 independent differentiation experiments. Each dot represents the mean area containing the EdU signal or the number of EdU puncta for one experiment of differentiation. The bar represents the mean of the 4 values, and the error bars show standard error of the mean. **(i)** Heatmap representing mRNA expression by RT-qPCR of genes encoding main actors of the UPR^mt^ at different time points (30 min, 2 h, 4 h, 6 h and 8 h) of mitochondrial stress, in the presence or absence of GSK2606414, compared to control conditions (DMSO only) at each time point. Data are represented in Log_2_(Fold change compared to Control) for 3 independent experiments (Type II Wald Chi-square tests ANOVA function with Tukey’s multiple comparisons test). mRNA expression was normalized relatively to *TBP* mRNA expression. *p-value ≤ 0,05; **p-value ≤ 0,01; ***p-value ≤ 0,001; **** p-value ≤ 0,0001

We next tested whether PERK inactivation could modulate stress-induced mitophagy (**Figure 4d, e**; **Supplementary Figure 3**). In the absence of antimycin A and oligomycin, there was no impact of GSK2606414 treatment on the number of neurons expressing PSer65-Ub. In contrast, application of GSK2606414 in stress conditions resulted in a notable decrease in the percentage of neurons expressing PSer65-Ub at all time points examined (from 60-65% with toxins only, to 38% with toxins and GSK2606414), demonstrating that induction of mitophagy upon mitochondrial stress was modulated by the PERK-ATF4 UPR^ER^. Strikingly, analysis of TOMM20 expression by immunofluorescence revealed that PERK inhibition by GSK2606414 resulted in a disorganization of the mitochondrial network in control conditions that was similar to that caused by mitochondrial stress (**Figure 4d, f**; **Supplementary Figure 3**). Adding GSK2606414 in stress conditions, however, did not trigger any changes in the spatial distribution of TOMM20 compared to the conditions with the toxins only or with GSK only. Moreover, GSK2606414 led to a strong alteration of mitochondrial biogenesis in control conditions (**Figure 4g, h**), as evaluated by the incorporation of EdU into de novo synthesized mtDNA molecules in control conditions or upon 8h of stress. Assessment of the number of EdU-positive puncta or the area of EdU+ signal per neuron revealed that PERK inactivation triggered a significant decrease in mitochondrial biogenesis in control conditions (90%) that was more drastic than that caused by treatment with the mitochondrial toxins only (60%); in contrast, GSK2606414 had no effect on mitochondrial biogenesis following stress. Thus, PERK-mediated UPR^ER^ appears to play a key role in maintaining the integrity of the mitochondrial network under basal conditions and in promoting mitophagy induction upon mitochondrial stress.

We then sought to determine whether the UPR^ER^ was responsible for attenuating the UPR^mt^ during mitochondrial stress (**Figure 4i**). We investigated mRNA expression of genes involved in the three UPR^mt^ branches and found that inhibition of PERK by GSK2606414 abolished or significantly reduced the stress-induced downregulation of *SIRT3* and the mitochondrial chaperones *CLPP*, *LONP1* and *YME1L1*, compared to control conditions. These results demonstrate a role for PERK in the inactivation of the ATF5- and SIRT3-mediated UPR^mt^ within 8 h of mitochondrial stress. In contrast, exposure to GSK2606414 reduced *NRF1* expression in control conditions from 30 min on, whereas it had no effect upon stress. In human DA neurons, PERK-mediated UPR^ER^ thus participates in regulating basal expression of *NRF1*, a key actor of the ERα-mediated UPR^mt^ response.

### Mitochondrial stress regulates the expression of LncRNAs in human DA neurons

One of the hypotheses raised to explain why midbrain DA neurons are more prone to degenerate in PD than other neuronal populations is that they may be particularly vulnerable to mitochondrial stress. In this context, we paid particular attention to lncRNAs, which constitute potent cell- and species-specific genomic regulators, speculating that they would be pivotal actors of the mitochondrial stress response specific to human DA neurons. From our transcriptomic data, we identified 1177 genes encoding lncRNAs expressed in human DA neurons (**Figure 5**). Amongst these non-coding elements, 23% had not been sequenced and annotated before and were therefore absent in existing databases (i.e. not annotated, **Figure 5a**). Using a categorization system based on their position relative to their closest protein-coding genes, we found that the majority of these lncRNAs were antisense overlapping (47%), intergenic (26%), bidirectional (12%) or sense overlapping (14%). Since most lncRNAs have not been functionally assessed yet, we estimated their putative functions considering their high probability to act in cis (Gil & Ulitsky, 2020) and thereby regulate their closest genes on the genome. Gene Ontology analysis on their adjacent protein-coding genes (**Figure 5b**) overall revealed a major enrichment in lncRNAs close to, and therefore potentially regulating, genes implicated in the regulation of transcription, highlighting the contribution of such elements in the regulation of genomic programs. We also found enrichments in genes involved in the regulation of developmental processes, telomere maintenance, or cytoplasmic translation. Among the 1177 lncRNAs, 336 were specifically expressed in the control condition and 159 only upon mitochondrial stress (**Figure 5c**). Interestingly, many lncRNAs with reduced expression or switched off upon stress were adjacent to protein-coding genes implicated in the regulation of transcription (**Figure 5d**), whereas many lncRNAs upregulated or specifically expressed under mitochondrial stress were at the vicinity of protein-coding genes involved in amino acid transport and regulation of nuclear division (**Figure 5d**). Furthermore, the proportion of novel, non-annotated lncRNAs, reached 28% within the stress-specific group, a proportion higher than among the control-specific lncRNAs (19%) or the lncRNAs expressed in both conditions (23%). Given the increasing number of existing and accessible RNA-sequencing data, this high percentage of newly discovered lncRNAs suggests a possible selective involvement in the response of human DA neurons to mitochondrial stress. Focusing on their closest protein-coding genes, we found an enrichment in terms associated with biological processes related to amino acid transport, as well as translation (**Figure 5e**). Thus, our data suggest that lncRNAs expressed upon mitochondrial stress contribute to the regulation of two major steps of the stress response of human DA neurons, as shown in **Figure 3b,d,f**. We next investigated whether lncRNAs expressed in DA neurons could be regulated by the transcription factors ATF3 and ATF4, which are the main mediators of the PERK UPR^ER^ (**Figure 5f**). Using available ChiP-seq datasets (Epanchintsev *et al*, 2017; Davis *et al*, 2018), we identified 571 putative binding loci for ATF3 and 202 for ATF4 within the promoters of 49% and 17% of the identified lncRNAs respectively. In both cases, we found that half of the potential target lncRNAs were regulated upon stress, around 40% of which were downregulated and 10% upregulated. The high proportion of lncRNAs potentially targeted for ATF3-dependent transcription, supports a preeminent role for the PERK-ATF4-mediated UPR^ER^ in the regulation of lncRNAs in human DA neurons exposed to mitochondrial stress. We selected lncRNAs of interest for further validation based on their expression profile upon stress, the function of their closest protein-coding genes and the presence of PD-associated single nucleotide polymorphisms (**Table 1**). We confirmed their expression profile by RT-qPCR and explored their regulation by the PERK-mediated UPR^ER^ using the inhibitor GSK2606414 (**Figure 5g**). In line with our previous results, expression of most lncRNAs was affected by PERK inhibition: GSK2606414 suppressed the stress-related downregulation of selected lncRNAs potentially involved in the generation and development of neurons (*lnc-TTC29*, *lnc-SLAIN1-11*, *lnc-MNAT1-2*, *ZNF778-DT*, *MIR4697HG*) and the upregulation of most of the selected lncRNAs associated with possible roles in the regulation of translation and the stress response (*lnc-SLC6A15, VLDLR-AS1, VPS11-DT, lnc-FKRP, SNHG1, TMEM161B-DT*). However, few lncRNAs were also regulated by GSK2606414 at basal level compared to control conditions (either downregulated, such as *FBXL19-AS1* and *NIPBL-DT*; or upregulated, such as *lnc-SLCA15-5* and *VPS11-DT*). Altogether, these results converge towards an implication of lncRNAs in the response of human DA neurons to mitochondrial stress, with a notable role in the regulation of translation mediated by the UPR^ER^.

**Figure 5.**
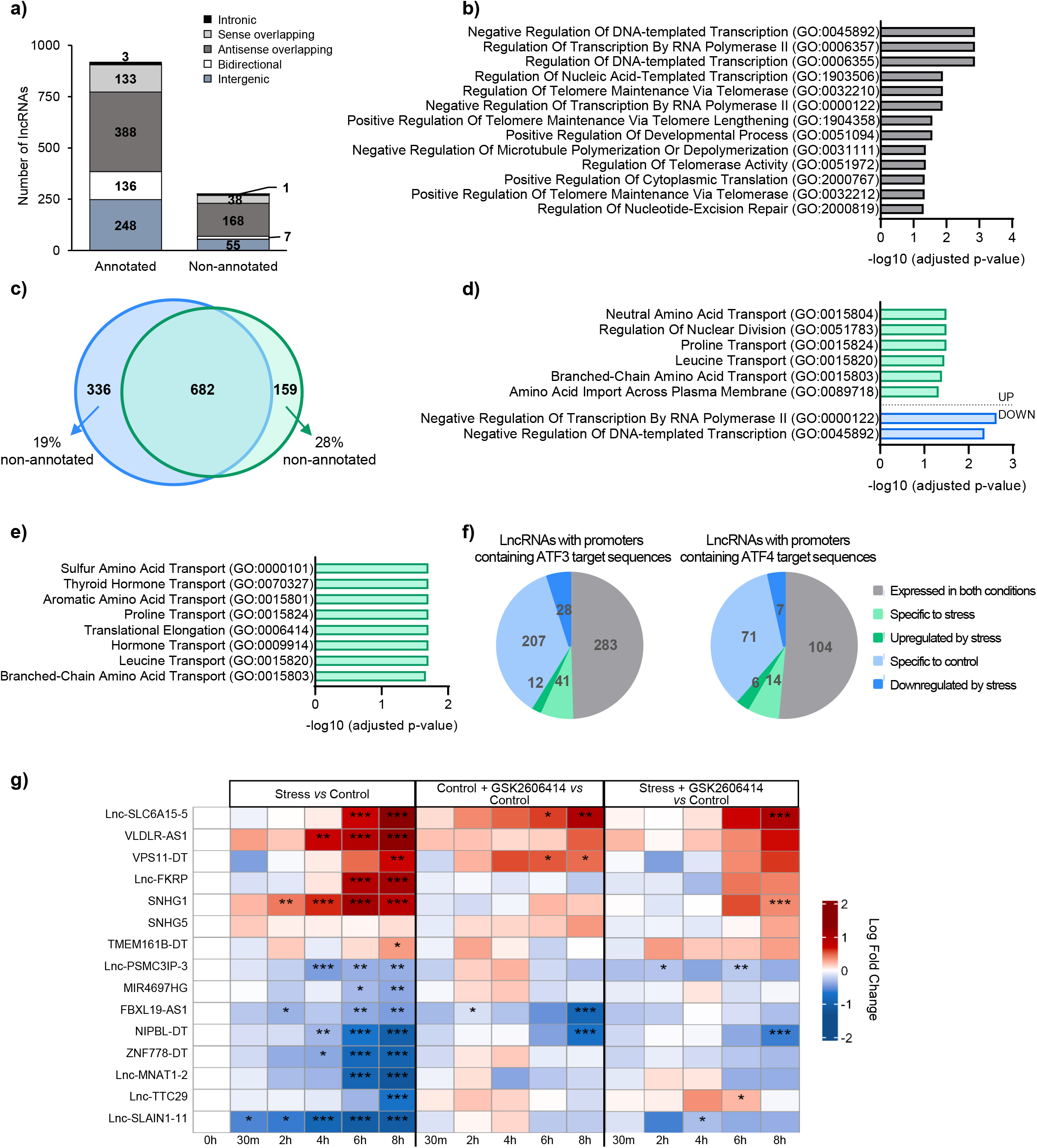
Mitochondrial stress response in human DA neurons includes the regulation of lncRNAs orchestrated by the PERK-mediated UPR^ER^. **(a)** Number of annotated and non-annotated lncRNAs depending on their genomic loci. n=4 RNA-seq datasets for DA neurons in control conditions; n=4 RNA-seq datasets for DA neurons submitted to a 8 h long mitochondrial stress. **(b)** Gene Ontology Analysis (Biological Process 2023, Enrichr) performed on the neighbouring protein-coding genes to all lncRNAs expressed in DA neurons, independent of the conditions (Control and Stress). **(c)** Venn diagram of overlap of lncRNAs expressed in control conditions and upon 8 h of mitochondrial stress. Percentages of lncRNAs expressed specifically in each condition are shown. **(d)** Gene Ontology Analysis (Biological Process 2023, Enrichr) performed on the neighbouring protein-coding genes to lncRNAs with altered expression upon stress (upregulated in green, downregulated in blue) compared to control conditions. **(e)** Gene Ontology Analysis (Biological Process 2023, Enrichr) performed on the neighbouring protein-coding genes to non-annotated lncRNAs specifically expressed upon stress. **(f)** Number of lncRNAs with promoters containing binding sites for ATF3 and ATF4 (determined by publicly available ChiP-Seq datasets) and categorized according to their expression profile upon stress compared to control conditions. **(g)** Heatmap representing RNA expression by RT-qPCR of genes encoding candidate lncRNAs (described in Table 1) at different time points (30 min, 2 h, 4 h, 6 h and 8 h) during mitochondrial stress, in the presence or absence of GSK2606414, compared to control conditions (DMSO only) at each time point. Data are represented in Log_2_(Fold change compared to Control) for 3 independent experiments (Two-way ANOVA with Tukey multiple comparisons test). RNA expression was normalized relatively to *TBP* mRNA expression. *p-value ≤ 0,05; **p-value ≤ 0,01; ***p-value ≤ 0,001

**Table 1.** Description of the selected lncRNAs. For each lncRNA presented in Figure 5g, the locus, classification depending on their genomic locus, closest coding-genes, presence of PD-associated single nucleotide polymorphism and presence of ATF3 or ATF4 binding sites at their promoter are provided.

| Category | Name | Locus | Classification | Closest protein-coding gene | Intersecting with PD-linked SNP | Promoter associated to ATF3 | Promoter associated to ATF4 |
| --- | --- | --- | --- | --- | --- | --- | --- |
| Upregulated in stress and associated to translation regulation and stress response | **Lnc-SLC6A15** | chr12:<br>83456731-84128270 | Intergenic | SLC6A15;<br>TMTC2 |  | Yes |  |
|  | **VLDLR-AS1** | chr9:<br>2503280-2621386 | antisense overlapping | VLDLR |  | Yes | Yes |
|  | **VPS11-DT** | chr11:<br>119067374-119067698 | bidirectional | VPS11 |  | Yes | Yes |
|  | **Inc-FKRP** | chr19:<br>46787815-46789039 | antisense overlapping | SLC1A5 |  | Yes |  |
|  | **SNHG1** | chr11:<br>62851991-62855460 | bidirectional | SLC3A2 |  | Yes | Yes |
|  | **SNHG5** | chr6:<br>85676990-85678736 | intergenic | SYNCRIP;<br>HTR1E |  | Yes | Yes |
|  | **TMEM161B-DT** | chr5:<br>88268938-88436684 | bidirectional | TMEM161B;<br>MEF2C |  | Yes | Yes |
| Downregulated and associated to PD-linked SNP | **Inc-PSMC3IP-3** | chr17:<br>42545451-42557693 | antisense overlapping | HSD17B1 | x |  |  |
|  | **MIR4697HG** | chr11:<br>133896438-133901601 | intergenic | IGSF9B;<br>SPATA19 | x |  |  |
|  | **FBXL19-AS1** | chr16:<br>30919319-30923269 | antisense overlapping | FBXL19 | x | Yes | Yes |
| Downregulated in stress and associated to neuron generation | **NIPBL-DT** | chr5:<br>36871364-36876700 | bidirectional | NIPBL |  | Yes |  |
|  | **ZNF778-DT** | chr16:<br>89215211-89217653 | bidirectional | ZNF778 |  | Yes | Yes |
|  | **Lnc-MNAT1-2** | chr14:<br>60657073-60659096 | antisense overlapping | SIX1 |  | Yes |  |
|  | **Inc-TTC29** | chr4:<br>146628898-146638145 | bidirectional | POU4F2 |  | Yes |  |
|  | **Inc-SLAIN1-11** | chr13:<br>78596129-78599619 | antisense overlapping | POU4F1 |  | Yes |  |

### The lncRNA *lnc-SLC6A15-5* specifically expressed in DA neurons regulates the resumption of translation following mitochondrial stress

We specifically focused on *lnc-SLC6A15-5*, a lncRNA selectively expressed upon mitochondrial stress (**Figure 6a**) and adjacent to the protein-coding genes *TMTC2*, involved in ER calcium homeostasis, and to *SLC6A15*, encoding a neutral amino acid transporter linked to depression, including in PD patients (Kohli *et al*, 2011; Zheng *et al*, 2017). Interestingly, *lnc-SLC6A15-5* was amongst the lncRNAs regulated by the PERK-mediated UPR^ER^ pathway at the basal level and following stress (**Figure 5g**). This lncRNA has been recently annotated (*ENSG00000289309*), but we identified 3 novel isoforms exclusively expressed in DA neurons exposed to mitochondrial stress (**Figure 6a, b**). These isoforms share the first 2 exons and the same transcription start site (TSS), at which level we observed an ATAC-seq peak that was significantly higher in the stress condition compared to control, suggesting the presence of an active promoter. The longest transcript possesses 5 exons, whereas the shortest isoforms display 3 and 4 exons respectively. Two additional annotated isoforms that were not sequenced in our datasets, *ENST00000689302.1* and *ENST00000688936.2*, have exons 2, 3 and 4 in common with the isoforms identified in our study, but have different TSS and first exon, not associated with an ATAC-seq peak. Similarly, another annotated lncRNA, *ENSG00000288941,* shared its last exon with the longest isoform detected here, but was not expressed in our datasets. Therefore, our newly annotated isoforms of *lnc-SLC6A15-5* may be specific to human DA neurons subjected to mitochondrial stress. To characterize further this lncRNA, we determined its subcellular localization using cellular fractionation, with *MALAT1* and *MT-ND2* as marker RNAs of the nucleus and the cytoplasm respectively and found that 65% of it was localized to the nucleus, whether or not the neurons were exposed to mitochondrial toxins (**Figure 6c**). We then sought to determine whether *lnc-SLC6A15-5* contributed to the major events of the mitochondrial stress response observed in human DA neurons: the alteration of the DA maturation status, the induction of mitophagy, the decrease in the de novo synthesis of mitochondrial DNA and the inhibition of general translation. We first used CRISPR inhibition technology coupled with viral vector delivery to knockdown this lncRNA. LUHMES cells transduced with a vector expressing either a single guide RNA targeting *lnc-SLC6A15-5* (sgRNA *lnc-SLC6A15-5)* or a sgRNA with no target (sgRNA NEG) were FACS-purified to generate homogeneous cell pools expressing each of these sgRNAs and differentiated into DA neurons. CRISPR inactivation led to an 85% decrease in *lnc-SLC6A15-5* expression upon stress compared to sgRNA NEG (**Supplementary Figure 4a**). Knock-down of *lnc-SLC6A15-5* had no effect on DA neuron identity or maturity, nor on induction of mitophagy or de novo synthesis of mitochondrial DNA at 8 h of stress (**Supplementary Figure 4b-g**). To investigate this lncRNA’s involvement in the regulation of translation following exposure to mitochondrial stress, we analyzed the incorporation of the puromycin analog O-propargyl-puromycin (OPP) into newly synthesized proteins in DA neurons with or without *lnc-SLC6A15-5* knockdown. As expected, mitochondrial stress applied for 8 h resulted in significant attenuation of translation compared to control conditions, as shown by the 81% decrease of the OPP signal (**Supplementary Figure 5a**), and in a significant increase of EIF2α phosphorylation (**Supplementary Figure 5b**). These effects were however independent of the changes in *lnc-SLC6A15-5* expression. We speculated that *lnc-SLC6A15-5* could be involved in the pro-survival response mediated by the resumption of translation after a stress. We therefore analyzed OPP incorporation 30 minutes after washing out the mitochondrial toxins from the culture medium to promote recovery of translation. There was no difference in levels of *lnc-SLC6A15-5* expression in the 30 minutes recovery condition compared to the mitochondrial stress condition without recovery (**Figure 7a**, NEG). Moreover, the knock-down approach reduced *lnc-SLC6A15-5* to about 15% of control levels in both conditions (**Figure 7a**, KD-Lnc-SLC6A15-5). Following wash out of the toxins from DA neurons treated with sgRNA NEG, there was a significant increase in the number of OPP puncta and intensity of OPP signal per neuron, indicating ongoing translational resumption (**Figure 7b-d, and Supplementary Figure 5**). This process was significantly slowed down in cells in which *lnc-SLC6A15-5* was knocked-down, as indicated by the reduction in OPP puncta and OPP signal intensity per neuron following removal of the toxins. These results suggest that *lnc-SLC6A15-5* contributes to a pro-survival response *via* the restoration of general translation. In this context, we explored the possibility that *lnc-SLC6A15-5* exerts this effect through inactivation of EIF2α (**Supplementary Figure 5c**) but did not detect any changes to the ratio between the phosphorylated form of EIF2α and total EIF2α levels following *lnc-SLC6A15-5* silencing. The UPR^ER^ triggers general translation attenuation not only *via* EIF2α but also through inhibition of mTOR. We therefore investigated the possible effect of *lnc-SLC6A15-5* on genes involved in the regulation of general translation *via* mTOR (**Figure 7e**, **Supplementary Figure 5d**). *SESN2* is a potent inhibitor of mTOR and a PERK-ATF4 target gene (Brüning *et al*, 2013; Garaeva *et al*, 2016). We found it to be overexpressed after stress in the KD-Lnc-SLC6A15 conditions compared to NEG *lnc-SLC6A15-5*, whether or not the toxins were washed out, indicating that it is regulated by *lnc-SLC6A15-5* under mitochondrial stress. Moreover, in the KD-Lnc-SLC6A15 condition, *SESN2* expression tended to increase more strongly following wash out of the toxins than in cells kept in presence of the toxins (p=0,09). Altogether, these results suggest that *lnc-SLC6A15-5* downregulated *SESN2* upon mitochondrial stress in human DA neurons, allowing for faster translation recovery once the stress signal becomes resolved. In parallel, we explored the role of *lnc-SLC6A15-5* in the regulation of genes encoding various amino-acid transporters (SLC1A3, SLC1A5, SLC3A2, SLC7A5) also known to be PERK-ATF4 target genes (Brüning *et al*, 2013; Han *et al*, 2013; Garaeva *et al*, 2016) and to contribute to mTOR activation (Zhuang *et al*, 2019). All these genes were found to be overexpressed following *lnc-SLC6A15-5* downregulation in cells exposed to mitochondrial toxins (**Figure 7e**), indicating a role of this lncRNA in their inhibition under conditions of mitochondrial stress. Strikingly, *SESN2* and the amino acid transporters regulated by *lnc-SLC6A15-5* are known PERK-ATF4 target genes (Brüning *et al*, 2013; Han *et al*, 2013; Garaeva *et al*, 2016), and are upregulated upon activation of the PERK-mediated UPR^ER^ (**Figure 3** Altogether, these results suggest that *lnc-SLC6A15-5* function counteracts ATF4-mediated transcription during mitochondrial stress. Accordingly, examining *ATF3* expression, we found it to be higher when a 30 minutes recovery from the toxin-induced stress was allowed in comparison to the full 8h of stress, in the condition of *lnc-SLC6A15-5* knock-down only (**Figure 7e**). This indicates an inhibitory effect of *lnc-SLC6A15-5* on *ATF3* expression during translation resumption after stress. Overall, *lnc-SLC6A15-5* appears to contribute to a pro-survival response associated with attenuation of PERK-ATF4-mediated UPR^ER^ and resumption of translation.

**Figure 6.**
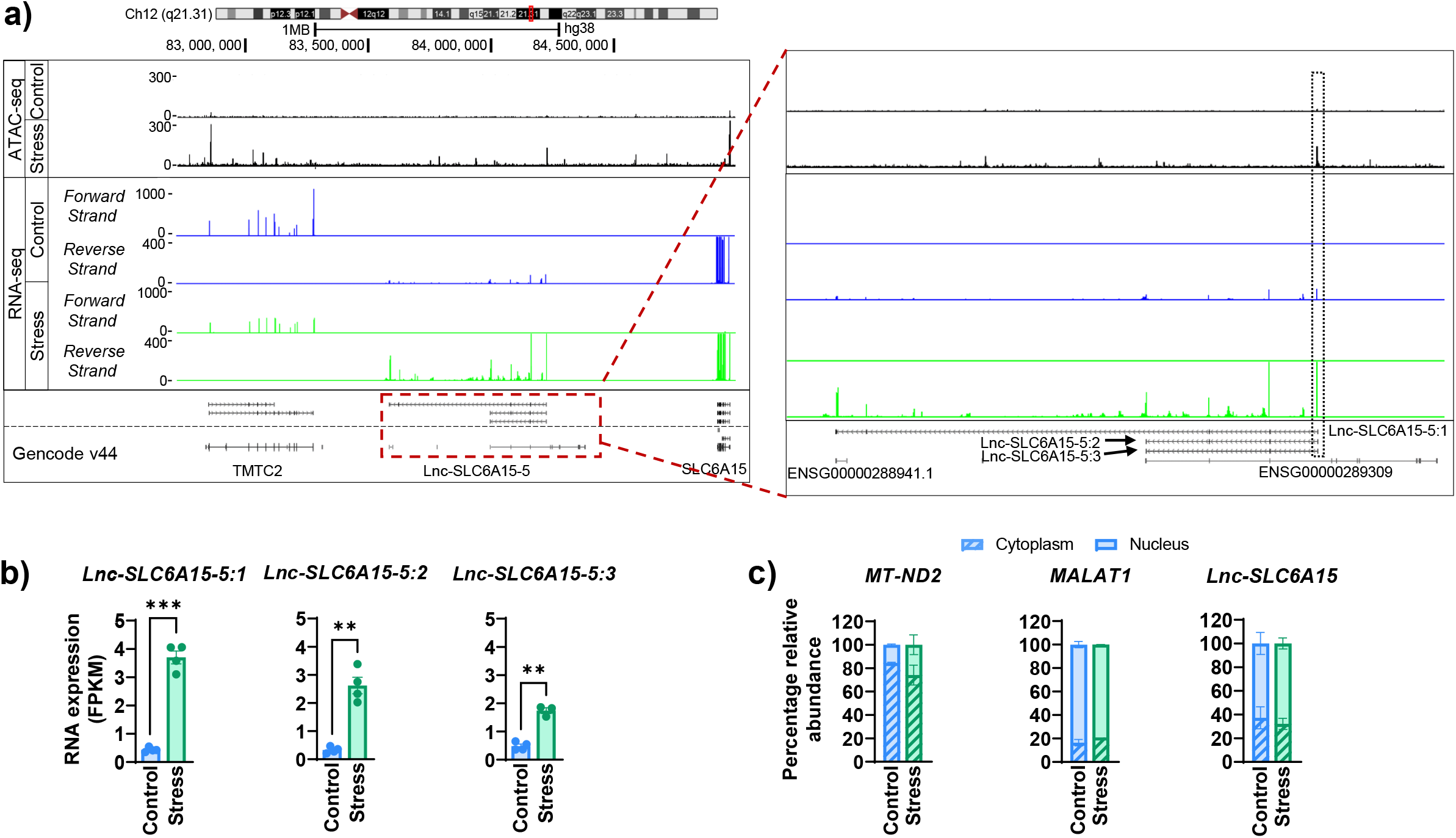
Newly identified *lnc-SLC6A15-5* is specifically expressed upon mitochondrial stress in DA neurons. **(a)** Schematics of the locus of *lnc-SLC6A15-5*. ATAC-seq peaks are depicted in black, reads from RNA-seq are in blue for the control condition, green for the stress condition. The scales represent reads per million (RPM). **(b)** RNA expression in FPKM of the 3 isoforms of *lnc-SLC6A15-5* in control (blue) or stress (green) conditions using the RNA-seq datasets (Control, n=4; Stress 8h, n=4). The bar represents the mean of the 4 values per condition (Control, Stress), and the error bars show standard error of the mean (DESeq2 differential analysis). Of note, transcripts are considered as expressed if their expression is higher than 1 FPKM in at least one sample and different from 0 in the 3 others. **(c)** Relative abundance of *lnc-SLC6A15-5*, assessed by RT-qPCR, in the nuclear or cytoplasmic fractions, in comparison with the nuclear RNA marker *MALAT1* and cytoplasmic RNA marker *MT-ND2*. Two independent experiments were used.

**Figure 7.**
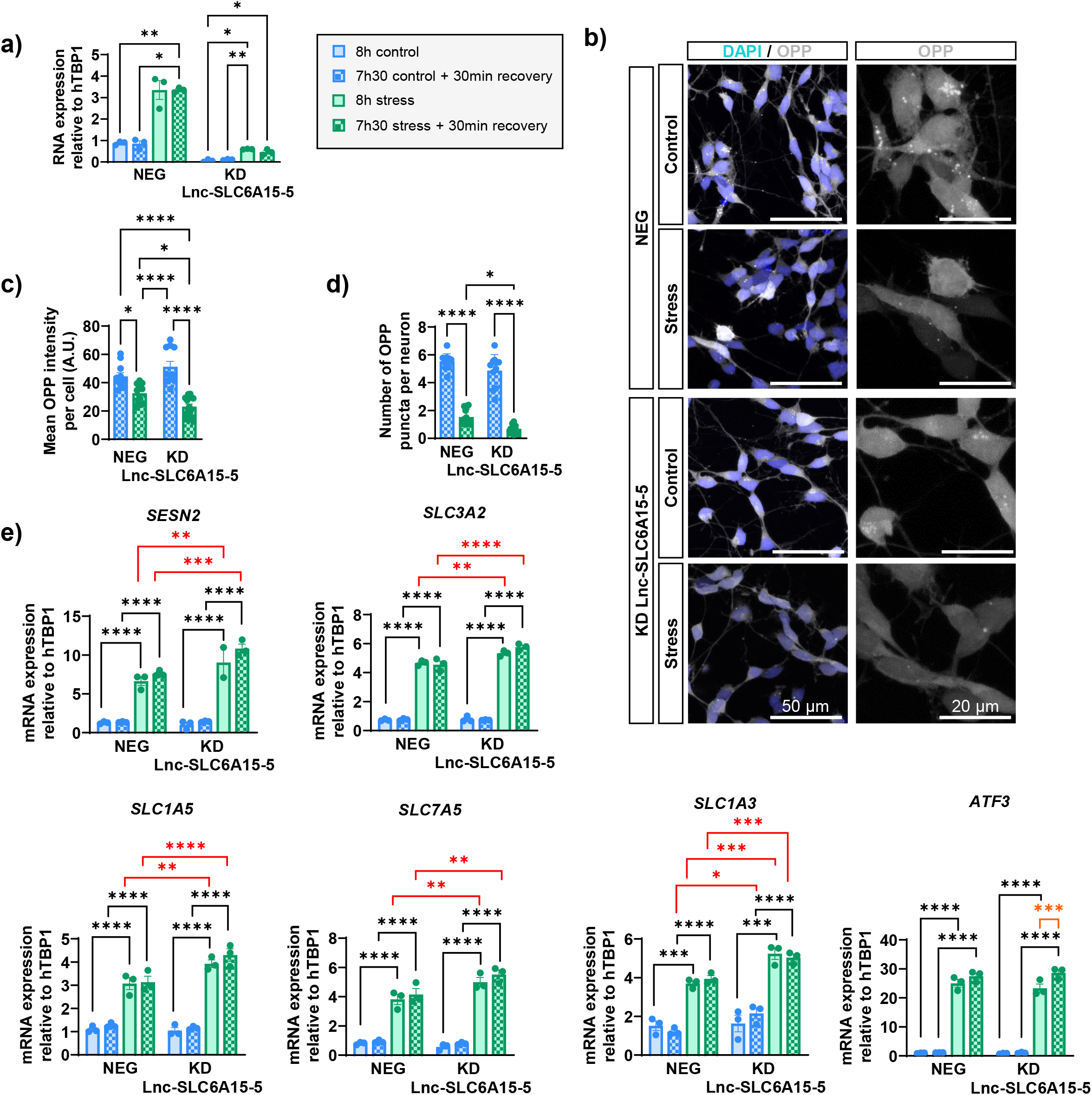
*lnc-SLC6A15-5* contributes to the resumption of translation following mitochondrial stress. **(a-e)** Four different experimental conditions were used. Control (blue) and stress (green) conditions were performed either for 8 h (plain bars) or for 7h30 followed by 30 min recovery (hatched bars). **(a)** *Lnc-SLC6A15-5* expression, assessed by RT-qPCR, in DA neurons transduced by lentiviral vectors carrying *dCAS9-KRAB* and sgRNAs either targeting lnc-SLC6A15-5 (KD *lnc-SLC6A15-5*) or a non-human sequence (NEG), in the 4 experimental conditions. RNA expression was normalized relatively to *TBP* mRNA expression. Data from 3 independent differentiation experiments, represented by 3 dots, were used. The bar represents the mean of the 3 values, and the error bars show standard error of the mean (Two-way ANOVA with Tukey’s multiple comparison test). **(b)** Detection of OPP (grey) in DA neurons expressing normal (NEG) or reduced levels (KD) of *lnc-SLC6A15-5*, in control or stress conditions for 7 h 30 min followed by 30 min recovery. Nuclei were stained using DAPI (blue). **(c)** OPP signal intensity in DA neurons (TH in control or stress conditions for 7 h 30 min followed by 30 min recovery (Two-way ANOVA with Tukey’s multiple comparison test). Data were obtained from 3 independent differentiation experiments. Each dot represents the percentage of the OPP intensity in DA neurons for one experiment of differentiation. **(d)** Number of OPP puncta per DA neuron, in control or stress conditions for 7 h 30 min followed by 30 min recovery (Two-way ANOVA with Tukey’s multiple comparison test). Data were obtained from 3 independent differentiation experiments. Each dot represents the number of OPP puncta per neuron per imaging field (n=4-6 per condition, per experiment). **(c and d)** The bar represents the mean of the values, and the error bars show standard error of the mean. **(e)** *SESN2, SLC1A3, SLC1A5, SLC3A2, SLC7A5 and ATF3* mRNA expression, assessed by RT-qPCR, in DA neurons expressing normal (NEG) or reduced levels (KD) of *lnc-SLC6A15-5*, in the 4 experimental conditions (Type II Wald Chi-square tests ANOVA function with Tukey’s multiple comparisons test). mRNA expression was normalized relatively to *TBP* mRNA expression. Data from 3 independent differentiation experiments, represented by 3 dots, were used. The bar represents the mean of the 3 values, and the error bars show standard error of the mean. *p-value ≤ 0,05; **p-value ≤ 0,01; ***p-value ≤ 0,001; **** p-value ≤ 0,0001

## Discussion

We conducted a comprehensive study to decipher and tie altogether the cellular and molecular mechanisms underlying the response of human DA neurons to mitochondrial stress. We demonstrated the central role of the PERK-mediated UPR^ER^in orchestrating cell-specific transcriptional programs upon stress, notably through the regulation of lncRNAs expression. Interestingly, PERK activation led to the inactivation of the UPR^mt^ and contributed to the maintenance of mitochondrial integrity and turnover following exposure to mitochondrial toxins. Importantly, we identified a stress-specific lncRNA, *lnc-SLC6A15-5*, which regulated the resumption of translation after mitochondrial stress, by modulating expression of ATF4 target genes involved in the regulation of mTOR activity.

Our work showed that inhibition of the electron transport chain by mitochondrial toxins triggered the concomitant activation of the 3 branches of the UPR^ER^ in human DA neurons, mediated by PERK, IRE1 and ATF6. However, the PERK-EIF2a-ATF4 pathway appeared to play a predominant role in the stress response, as its pharmacological inhibition resulted in alterations at the transcriptional and cellular levels that counteracted this response. While there is a consensus regarding the importance of ATF4 in the mitochondrial stress response in mammalian cells, the involvement of the other branches of the UPR^ER^ or UPR^mt^ remains unclear. The fact that many crosstalks have been uncovered between the PERK-, IRE1- and ATF6-mediated UPR^ER^ (Walter *et al*, 2018; Brewer, 2014) makes the analysis of the contribution of each pathway in the stress response more complex. Indeed, PERK-ATF4 involvement could mask activation of other pathways that share common target genes. Moreover, despite the use of cellular models with homogeneous cell populations, such as LUHMES cells, stress responses differ depending on the cell types, the activating stimuli, or the duration and intensity of the mitochondrial stress signal (Lamech & Haynes, 2015; Ko *et al*, 2020). This is particularly noteworthy regarding the activation of the UPR^mt^, which has been observed more frequently, but not systematically, after longer stressor applications (Houtkooper *et al*, 2013; Krug *et al*, 2014; Monti *et al*, 2015; Quirós *et al*, 2017; Cai *et al*, 2020), or using stressors leading to localized alterations, such as mitochondrial protein folding stress (Münch & Harper, 2016; Uoselis *et al*, 2023). In our study, we found that the UPR^mt^ was inactivated following 8 h of treatment with the chosen mitochondrial toxins, and remarkably this effect was directly attributable to the PERK-UPR^ER^. We cannot exclude that we only observed the first steps of a multiphasic stress response in which UPR^mt^ would be induced at later time points. However, our results suggest that the 8 h stress protocol brought the cells to a crossroad where pro-survival, protective and pro-apoptotic pathways were co-activated, but ultimately leaning towards commitment to cell death, as indicated by the increase in apoptotic neurons observed in our conditions and the predominant activation of the PERK-UPR^ER^ with upregulation of the pro-apoptotic factors CHOP, CHAC1 and TRIB3.

At the cellular level, we have shown that the PERK-UPR^ER^ also contributes to the regulation of mitochondrial turnover upon mitochondrial stress in human DA neurons, since its inactivation by the PERK-specific pharmacological inhibitor GSK2606414 significantly reduced the proportion of DA neurons with ongoing mitophagy. Such direct implication of PERK activation in the induction of mitophagy has been sparsely documented so far and only been described following exposure to chromium (Dlamini *et al*, 2021) or ER stress (Zhang *et al*, 2014), through the transcriptional activation of *PRKN* expression by ATF4 (Zhang *et al*, 2014; Bouman *et al*, 2011). In contrast, ATF4 appeared to restrain the induction of mitophagy following mitochondrial protein folding stress (Uoselis *et al*, 2023). In our study, *PRKN* was not regulated transcriptionally upon stress, suggesting that modulation of mitophagy by PERK-ATF4 could be channeled through another mechanism.

In contrast, activation of PERK was not found to contribute to the changes in mitochondrial biogenesis, monitored upon stress in terms of newly synthesized mitochondrial DNA. However, using the inhibitor GSK2606414, we showed that PERK pathway activity was necessary in basal conditions for the maintenance of the mitochondrial network and for the biogenesis of mitochondrial DNA, in line with the literature (Muñoz *et al*, 2013; Mesbah Moosavi & Hood, 2017; Kato *et al*, 2020; Sassano *et al*, 2023). Remarkably, these studies revealed that PERK involvement was independent of the UPR. In parallel, ATF6 has also been linked to mitochondrial biogenesis, namely through its regulation by PGC1α (Wu *et al*, 2011; Misra *et al*, 2013). In our conditions, involvement of PERK was demonstrated *via* its pharmacological inactivation, which completely shut down mitochondrial DNA biogenesis. However, we cannot rule out crosstalk between PERK- and ATF6-mediated pathways, with PERK being the primary sensor.

To assess the intrinsic features governing the cell-specificity of the response of human DA neurons to mitochondrial stress, we focused on lncRNAs. Expression of such molecules only depends on transcription, and therefore they can be rapidly mobilized in the context of the cellular response to stress. In addition, their regulatory functions in crucial cellular processes, such as cell growth, proliferation, apoptosis or translation, as well as their species- and cell-specificity, constitute strong arguments for a role of lncRNAs in adapting stress responses to the particularities of each cellular subtypes. LncRNAs have been shown to participate to regulatory pathways associated with p53, mTOR and eIF2 (Scholda *et al*, 2023). A growing number of studies have explored the contribution of lncRNAs to ER stress response (Quan *et al*, 2018; Li *et al*, 2023), and very often in the context of pathologies such as cancers, leading to the identification of disease-related molecular signatures (Zhang *et al*, 2023; Chen *et al*, 2022; Shen *et al*, 2023). However, most of these investigations focused on single lncRNAs, many of which directly regulate components of the UPR^ER^ (Brookheart *et al*, 2009; Yang *et al*, 2015; Bhattacharyya & Vrati, 2015; Su *et al*, 2016; Wu *et al*, 2020; Martinez-Amaro *et al*, 2023). In line with this, we have discovered novel isoforms of a lncRNA, *lnc-SLC6A15-5*, with probable roles in the resumption of translation once the mitochondrial stress is resolved. Interestingly *lnc-SLC6A15-5* appeared to contribute to the regulation of the UPR^ER^, as it exerted its action on the transcription of ATF4 target genes that encode mTOR modulators. *Lnc-SLC6A15-5* was upregulated by the UPR^ER^ upon stress, overall suggesting that it could be part of a feedback loop to dampen the UPR^ER^ once the stress is over.

So far, genome-wide studies investigating lncRNAs in the context of PD relied on existing databases, and most of them were performed on brain tissue or blood samples from patients or animal models (Xin & Liu, 2021). Such approaches bring important information regarding lncRNAs as potential biomarkers of the disease, however they have limited potential for the discovery of cell-specific lncRNAs and their functions (Liu *et al*, 2017; Mattick *et al*, 2023). Thus, the vast majority of the lncRNAs investigated in PD are ubiquitously expressed and also known for their implication in other diseases, including MALAT1, NEAT1, H19, lncRNA-p21 or SNHG1 for instance (Kraus *et al*, 2017; Yan *et al*, 2018; Qian *et al*, 2019; Liu *et al*, 2020; Zhang *et al*, 2020; Xin & Liu, 2021; Zhang *et al*, 2022). Using a method allowing for the discovery of novel transcripts, we have established the exhaustive repertoire of lncRNAs expressed in mature human LUHMES-derived DA neurons in basal conditions and following mitochondrial stress. This study lays the foundations for the detailed investigation of the role of lncRNAs in key steps of the DA-specific response to mitochondrial stress and, more generally, in the pathophysiology of PD, characterized by the degeneration of these neurons.

## Materials and Methods

### LUHMES cell culture and differentiation

LUHMES cells were grown in proliferation medium containing Advanced Dulbecco’s Modified Eagle Medium (DMEM)/F12, 1% N-2 supplement, 1% penicillin/streptomycin (P/S), L-Glutamine (2 mM, Life Technologies), and human basic fibroblast growth factor (FGF, 40 ng/mL, R&D Systems). Cells were maintained at 37°C in a humidified atmosphere containing 5% CO_2_, passaged using 0.05% trypsin (Gibco) and plated at a density of 2.3 x 10^4^ cells/cm^2^. Plastic cell culture flasks and multi-well plates were coated with poly-L-ornithine (pLO, 50 μg/mL), fibronectin bovine plasma (1 μg/mL, Sigma-Aldrich) and 1% P/S, and incubated overnight at 37°C. After removing the coating solution, culture flasks were washed twice with water before cell seeding.

For differentiation, LUHMES cells were plated at a cell density of 5 x 10^4^ cells/cm^2^ in proliferation medium. After 24 h (day 0 of differentiation), proliferation medium was replaced by differentiation medium consisting of Advanced DMEM/F12, 1% N-2 supplement, 1% P/S, L-Glutamine (2 mM), dibutyryl cyclic AMP (cAMP, 1 mM, Sigma-Aldrich), recombinant human growth-derived neurotrophic factor (GDNF, 2 ng/mL, Peprotech) and tetracycline (1 μg/mL, Sigma-Aldrich). On day 2 of differentiation, LUHMES cells were seeded into pre-coated culture plates at a cell density of 1x 10^5^ cells/cm^2^. The following day, differentiation medium was changed.

### Mitochondrial Stress

LUHMES-derived DA neurons (day 6 of differentiation) were treated with a combination of antimycin A (25 µM, Sigma-Aldrich), and oligomycin (10µM, Sigma-Aldrich). Stock solutions of these toxins, at 2 mg/mL and 25 mg/mL respectively, were dissolved in dimethyl sulfoxide (DMSO). After treatment, neurons were collected or fixed for subsequent analysis. For controls experiments, DMSO was added to the samples without mitochondrial toxins.

### PERK-UPR^ER^ inhibition

For experiments investigating the contribution of the PERK-mediated UPR^ER^ to the stress response, cells were incubated with GSK2606414 (25µM, Selleckchem), simultaneously to the incubation with mitochondrial toxins or with DMSO for the control experiments.

### Immunofluorescence

Glass coverslips were added to the 4-well plates and pre-coated with pLO and fibronectin, overnight at 37°C. Next, laminin (5 μg/mL, Life technologies) was added to the coating medium at 37°C for 1 h, before cell seeding. After culture and treatments, cells were fixed for 15 min in 4% paraformaldehyde prepared in PBS 1X. Immunofluorescent labelling was performed as described previously (Gendron *et al*, 2019). The primary and secondary antibodies used are described in Supplementary Table S1. Nuclei were labelled with DAPI DNA stain.

Image acquisition were performed on either SP8 inverted confocal microscope (Leica) with a 40x or 63x oil immersion objective or AxioScan Z1 (Zeiss) with a x20 objective.

### Mitochondrial biogenesis assay

Replication of mitochondrial DNA (mtDNA) was monitored using the Click-iT Plus EdU Cell Proliferation Kit for Imaging (Salic & Mitchison, 2008) (Invitrogen) according to manufacturer’s instructions. Briefly, EdU was added at a final concentration of 10 μM to the differentiation medium, and incubated for 2, 4, and 8 h at 37°C, or only 8 h when cells were treated with GSK2606414. Then, neurons were fixed in 4% PFA for 15 min at room temperature, followed by permeabilization with PBS-Triton 0.2% supplemented with 4% goat serum overnight at 4°C. The following day, after washes, cells were incubated for 30 min at room temperature in the dark with the Click-iT Plus reaction cocktail readily prepared according to the manufacturer’s instructions. In the EdU control experiments, neurons were pretreated for 4 h with 2’,3’-dideoxycytidine (ddC, 100 μM). This was followed by cotreatment of ddC and EdU for 4 h or 24 h of incubation. As ddC was dissolved in DMSO, controls experiments were supplemented with an equal DMSO volume. For control experiments regarding mitochondrial stress, DMSO was added to the samples without mitochondrial toxins.

### Translation assay

Protein synthesis was assessed using the Click-iT Plus OPP Alexa Fluor 647 Protein Synthesis Assay Kit (Invitrogen) according to manufacturer’s guidelines. Cells were incubated in fresh medium containing O-propargyl-puromycin (OPP, 20 µM) for 30 min at 37°C. For this experiment, two conditions were tested: 8 h stress or DMSO for which the OPP-supplemented medium also contained mitochondrial toxins or DMSO, and 7h30 stress/DMSO followed by 30 min of recovery for which the OPP-supplemented medium was free of toxins or DMSO. After 30 min of incubation, cells were fixed using 4% PFA for 15 min at room temperature and, after washes in PBS1X, stored at 4°C overnight in PBS1X. The following day, cells were permeabilized in PBS-Triton 0.5% for 15 min and were then incubated for 30 min at room temperature in the dark with freshly made Click-iT OPP reaction cocktail as per manufacturer’s instructions. Cells were then washed with the Click-iT Reaction Rinse Buffer before proceeding with immunofluorescence and DAPI staining.

### RNA extraction

Total RNAs were purified from LUHMES cells using a RNeasy Minikit (Qiagen) following manufacturer’s instructions. RNAs were further treated with DNAse I (Roche) for 20 min at room temperature to prevent genomic DNA contamination. RNA quantification was determined either by spectrophotometry (Nanodrop 2000c, THERMO Scientific) prior to RT-qPCR or using a High Sensitivity RNA ScreenTAPE analyzer (Agilent technologies) for RNA-seq. In the latter case, the RNA integrity number (RIN) was used to determine RNA quality for all tested samples. RNA was stored at −80 °C until reverse transcription or RNA-seq.

For subcellular localization, Trizol reagent (Life technologies) was used following manufacturer’s instructions for RNA extraction.

### Subcellular fractionation

A minimum of 10 million of cells was used for subcellular fractionation. LUHMES cells were enzymatically dissociated by using 0.05% trypsin, centrifuged, and washed once with PBS. The cell pool was divided in two parts, the first part for total RNA extraction directly lysed with Trizol reagent (Life technologies) and -80°C frozen, and the second part for fractionation. Subcellular fractionation was performed as described by Gagnon et al. (Gagnon *et al*, 2014). Briefly, cells were lysed with hypotonic buffer (10 mM Tris-HCl [pH 7.5], 10mM NaCl, 3mM MgCl_2_, 0.3% NP-40, 10% Glycerol) supplemented of 100U RNase-OUT and DTT (10mM) during 10 min on ice. Intact nuclei were separated of cytosol fraction by centrifugation 3 min at 1000g at 4°C. Then, the supernatant was recovered (cytosol fraction), and RNA were precipitated with 150mM Na_2_Ac [pH 5.5], 95% EtOH supplemented with DT40 (10µg), 1 h at -20°C. During this time, the pellet was washed in hypotonic buffer (see above) before lysed by Trizol reagent supplemented with DT40 (10µg) and then frozen at - 80°C.

### Real time quantitative RT-PCR (RT-qPCR)

Up to 800 ng total RNAs were used to generate a first cDNA strand (Superscript II reverse transcriptase, THERMO Fisher Scientific) with random hexamers as indicated by the manufacturer. qPCR experiments were realized on the Light Cycler 384 real-time PCR system (Roche); with SYBER green detection (Roche). The comparative method of relative quantification (2^−ΔΔCT^) was used to calculate the expression levels of each target gene and human TBP mRNA was used to normalize the expression of all samples. The list of primers used is provided in the Supplementary table S2.

### Western blotting

Cells were differentiated and re-plated in T75 flasks. At day 6 of the differentiation, cells were scraped from the plate and lysed in 1× Ripa buffer (Sigma-Aldrich) supplemented with 1× protease and phosphatase inhibitor cocktail (ThermoFisher Scientific). The lysate was centrifuged at 13 500 rpm for 5 min at 4 °C, and the supernatant was collected. Protein concentration was measured using a BCA kit (Pierce). Each sample (15 μg) was boiled for 5 min and applied on NuPAGE 4%– 12% Bis-Tris Gel (Biorad). The gel was transferred onto a nitrocellulose membrane. The membranes were incubated with primary antibodies at 4°C overnight followed by secondary antibody for 1 h at room temperature. Membranes were scanned and analyzed using Chemidoc Touch Imaging system (Biorad). The list of antibodies used for western Blotting are presented in Supplementary Table S1.

### Image processing and analysis

For experiments investigating mitophagy induction, the slides were digitized using the AxioScan Z1 (Zeiss) with a x20 objective and acquired using the ZEN software. The resulting files were exported and whole slide image were processed on QuPath, an open-source machine learning software (Bankhead *et al*, 2017). The total number of cells on each image was obtained by counting the DAPI stained nuclei. Following cell detection, the QuPath algorithm was able to quantify the number of cells labeled with markers of interest by setting specific intensity thresholds. For experiments investigating mitophagy, mitochondrial biogenesis and protein synthesis, images were acquired with the Leica TCS SP8 Digital LightSheet inverted confocal microscope with a x40 or x63 oil objective, using the LAS (Leica Application Suite) X acquisition software and processed with the ImageJ software available at https://imagej.nih.gov/ij/ (Schneider *et al*, 2012). A threshold was set to select the signal of interest, then different parameters were analyzed including the area and intensity of our signal of interest (here, EdU and OPP). For punctiform signals, as provided by EdU and OPP, the number of puncta per neuron (and puncta colocalization with TOMM20 for EdU) was also quantified using the spot detection and colocalization ImageJ plugin ComDet v.0.5.5. Regarding the analysis of the mitochondrial marker TOMM20 labeling, the area occupied by clusters of mitochondria (defined as the area of TOMM20+ clusters) was also measured using the subcellular detection tool from QuPath software.

### DNA lentiviral constructs for CRISPR inhibition and activation

Loss of function was performed with CRISPR inhibition technology (CRISPRi). LV_U6-empty_EF-1α-KRAB-dCas9-T2A-TagEGFP backbone vector was kindly provided by Jorge FERRER’s Lab (Imperial College London). The sgRNA targeting *lnc-SLC6A15-5* (KD), or the control sgRNA (sgNEG - targeting the human AAVS1 locus), were cloned into BsmBI sites of LV backbone vectors. The KD sgRNA was designed 35pb after *lnc-SLC6A15-5* transcription start site (TSS) using a bulge-allowed quick guide-RNA designer for CRISPR/Cas derived RGENs (http://www.rgenome.net/cas-designer/).

All sgRNA sequences used in this study are presented in Supplementary Table S3.

### Lentiviral vector production

Lentiviral vector stocks were produced as previously described (Scharfmann *et al*, 2008). Briefly, HEK 293T cells were transfected by the p8.9 packaging plasmid (ΔVprΔVifΔVpuΔNef) (2), the pHCMV-G that encoded the VSV glycoprotein-G (Zufferey *et al*, 1997) and the pTRIP ΔU3 recombinant lentiviral vector. The supernatants were treated with DNAse I (Roche Diagnostic) prior to their ultracentrifugation, and the resultant pellets were re-suspended in PBS, aliquoted, and then frozen at -80°C until use. The amount of p24 capsid protein was quantified by the HIV-1 p24 antigen ELISA (Beckman Coulter). All transductions were normalized relative to p24 capsid protein quantification.

### Fluorescence-activated cell sorting (FACS)

For CRISPRi experiments, KD-lnc-SLC6A15-5 or control (NEG) cells are both expressing GFP and were purified using cell sorting. Cells were enzymatically dissociated by using 0.05% trypsin, centrifuged, washed once with PBS and filtered (50 μm filter) prior to cell sorting. GFP+ cells were purified using a S3 Biorad cell sorter. Cell suspensions from LUHMES non-transduced were used to adjust background fluorescence.

### RNA-sequencing (RNA-seq)

4 independent LUHMES cell differentiations were treated or not with mitochondrial toxins. 500 ng of total RNA were used from Control (n=4) and stressed (n=4) DA neurons to prepare stranded RNAseq libraries following manufacturer’s recommendations using KAPA mRNA hyperprep (Roche Diagnostic). Each final library was quantified and qualified with 2200 Tapestation (Agilent). Final samples of pooled library preparation were sequenced on NextSeq500 with High Output Kit cartridge at 2×150M reads/sample.

### Assay for Transposase-Accessible Chromatin (ATAC-seq)

For each sample, 70 000 cells collected and centrifuged at 500 g, at 4 °C during 20 min. Cells were resuspended in 25 μL of lysis buffer (10 mM Tris-HCl pH 7.4, 10 mM NaCl, 3 mM MgCl_2_-6H_2_O, 0.1% IGEPAL CA-630) during 30 min at 4 °C. Then, after centrifugation at 500 g, at 4 °C during 30 min the nuclear pellet was treated by Tn5 transposase. The pellet was resuspended in 25 μl of 12.5 μl 2x TN buffer; 2 μl of Tn5; 10.5 μl d’H_2_O and incubated at 37 °C for 1 h. Next, 5 μl of clean-up buffer (900 mM NaCl, 300 mM EDTA) were added to transposase treated nuclei, followed by 2 μl of 5% SDS and 2 μl of 20mg/ml Proteinase K, and incubated for 30 min at 40 °C. DNA samples were then purified twice using 68 μl of AMPure-XP beads (Beckman Coulter_A63881) and next eluted in 13 μl of buffer EB (Qiagen Cat No./ID: 19086). Amplification and size selection of ATAC-seq libraries were performed according to Grbesa et al. (2017 PMID: 29155775) using Nextera XT Index kit (Illumina-15055293). Extracted DNA concentration was measured by 2200 Tapestation (Agilent Technologies). Final samples of pooled library preparation were sequenced on Novaseq6000 with SP-100 cartridge at 100M reads/sample.

### RNAseq analysis and de novo annotation of lncRNAs

The analysis of lncRNA expressions from Next-Generation Sequencing (NGS) data involved a comprehensive pipeline of sequential steps. Initially, raw FASTQ files underwent quality assessment using FastQC v0.11.8, followed by Trimmomatic v0.39 trimming to remove low-quality trailing bases, adapters, and reads shorter than 50 bases. Cleaned reads were then aligned to the hg38 human reference genome using HISAT2 v2.2.1 (Kim *et al*, 2019), resulting in ordered BAM files generated through Samtools v1.11 (Danecek *et al*, 2021). Subsequent transcript assembly and abundance estimation were performed using StringTie v2.1.4 (Pertea *et al*, 2015), followed by the merging of transcript annotations from all samples into a unified catalog using StringTie merge. Expression levels of transcripts were quantified through StringTie FPKM normalization. Comparative analysis against Gencode v44 and LNCipedia v5.2 reference catalogs was carried out using GffCompare v0.11.2 (Pertea & Pertea, 2020), with transcript annotations categorized as “known” or “unknown” based on class codes. Coding potential prediction was executed using CPC2 v1.0.1 (Kang *et al*, 2017), CPAT v3.0.3 (Wang *et al*, 2013) and CNIT (Guo *et al*, 2019). Annotations were enriched with details about nearest protein-coding genes and LNCipedia classification. The catalog underwent successive filtration, including removal of low-expression transcripts, retention of non-coding transcripts predicted by multiple tools, and elimination of short transcripts with lengths below 200 bases. Additionally mono-exonic transcripts not present in Gencode or LNCipedia were retained only if an ATAC-seq peak was present within 100 bases from the transcription start site (TSS). The remaining transcripts were filtered according to specific gene/transcript types from Gencode. The filtered catalog was then merged with the Gencode catalog, appending transcripts that did not exactly match the reference. This updated catalog was quantified with STAR v2.7 (Dobin *et al*, 2013) using original FASTQ files. Resulting FPKM counts were integrated into the filtered catalog, which underwent consolidation into a gene-centric format, retaining annotations solely for the most highly expressed transcript per gene. This comprehensive pipeline facilitated the detailed analysis of lncRNA expressions and provided valuable insights into their roles and functions. Normalization and differential analysis for protein-coding or non-coding genes were performed with the DESeq2 package.

### ATAC-seq data processing

Steps for quality control were identical to those used for RNA-seq data treatment (Trimmomatic, FastQC). Reads with a length below 50 bp have been removed in further analysis. Paired-end reads were mapped to the human genome (build hg38) with Bowtie2. Duplicate reads were discarded with the Picard tools. Peaks were called using the MACS2 program with the option callpeak. Individual peak annotations were obtained with the R software version 3.5.1 (R Development Core Team, 2018) using the ChIPseeker R package (v1.20). Consensus peak was obtained using the DiffBind R package (v2.12).

### Pathway enrichment analysis and transcription factor motifs search

Enrichr web tool (https://maayanlab.cloud/Enrichr/) was used to perform gene ontology (GO) and pathway enrichment analysis of gene lists with the GO Biological Processes 2023 and Reactome 2022 databases (Chen *et al*, 2013; Kuleshov *et al*, 2016; Xie *et al*, 2021; Gillespie *et al*, 2022). Cistrome Data Browser toolkit (http://dbtoolkit.cistrome.org) and Cistrome-GO were used to identify transcription factors with binding sites significantly overlapping promoters of lncRNAs or ATAC-seq-detected open chromatin regions, and to perform functional enrichment analysis from the results obtained (Zheng *et al*, 2019; Mei *et al*, 2017; Li *et al*, 2019).

### Data availability

Raw sequence reads from RNA-seq and ATAC-seq are available from GEO under accession number GSE (in progress).

### Statistics

For evaluation of TH, DAT and cCASP3 staining as well as P-eIF2α/eIF2α protein levels in 8 h control and stress conditions, statistical analyses were performed using paired Student’s *t* tests. For mitochondrial DNA synthesis, mitophagy and protein synthesis experiments, two-way analysis of variance ANOVA tests were applied, followed by post-hoc Tukey’s multiple comparison test. All these tests were carried out using GraphPad 9.1.2. Regarding kinetic experiments evaluating RNA expression of UPR^ER^ and UPR^mt^ factors as well as of candidate lncRNAs, group differences and evolutions of expression values were investigated using linear mixed-effects models (LMMs) by fitting one model per gene of interest. In each model, the factor variables Condition (four levels for Control and Stress with or without the GSK2606414 treatment), Time (30, 120, 240, 360, and 480 minutes) and their interaction term were regarded as fixed effects, while a random (intercept) effect was used to account for values obtained from the same differentiation experiments. All LLMs were fitted using the lmer function of the lme4 R package (v1.1-34) (Bates *et al*, 2015) with R version 4.3.1 (R Development Core Team, 2023). For each gene, the significance of the main and interaction effects of Condition and Time was assessed by Type II Wald Chi-square tests using the function Anova of the car R package (v3.1-2). For post hoc pairwise comparisons, all conditions were compared at each time point using the emmeans R package (v1.8.8) with the Tukey’s method for multiple testing. Prior to modeling, a log-transformation (log(x+0.1)) was applied to expression data in order to better meet the the LMM assumptions of normality and homoscedasticity of residuals. The same analysis was performed on the RT-qPCR results looking at the RNA expression of multiple targets in the 7h30 with 30 minutes recovery or 8 h control or stress conditions. All the test results were graphically reported as heatmaps generated with the ComplexHeatmap R package (v.2.16.0).

### Supplemental material

Results from additional experiments are shown in Supplementary Figures 1 to 5: Supplementary Figure 1 shows the incorporation of EdU in mtDNA in DA neurons as mean to follow mitochondrial biogenesis. Supplementary Figure 2 shows the number of ATAC-seq peaks defining chromatin regions with altered accessibility upon 8 h of mitochondrial stress, as well as Gene Ontology enrichment analyses performed on genes associated with these regions. Supplementary Figure 3 shows the effect of GSK2606414 on the initiation of mitophagy (marked by the expression of Phospho-Serine 65 ubiquitin) following 4 and 6 h of mitochondrial stress, compared to control conditions. Supplementary Figure 4 shows the effect of the inhibition of Lnc-SLC6A15-5 expression in the number of mature DA neurons (TH and DAT expression *via* immunofluorescence), initiation of mitophagy (Phospho-Serine 65 ubiquitin expression) and mitochondrial biogenesis (Edu integration into mtDNA) upon 8 h of stress. Supplementary Figure 5 shows the effect of the inhibition of Lnc-SLC6A15-5 expression on the expression of molecular actors involved in the regulation of translation, at the protein level (EIF2α and phospho-EIF2α) or mRNA level (DDIT4, SYNCRIP, MTOR, RPS6KB1, RPS6, EIF4EBP1 and EIF4EBP2), following 8 h of stress, or 7h30 of stress and 30 min of recovery. Supplementary tables 1, 2 and 3 recapitulate the lists of antibodies, primer sequences and sgRNA sequences used in this study.

## Data availability statement

The GEO accession number for RNA-seq and ATAC-seq reported in this paper is: All other data are available in the manuscript or supplemental materials.

## Acknowledgments

The authors would like to thank the iVECTOR core facility of the Paris Brain Institute for technical assistance in producing all lentiviral vectors, the Genotyping and Sequencing Platform of the Paris Brain Institute for technical assistance in performing RNA-seq, and ATAC-seq. We thank Jorge Ferrer (Imperial College London) for sharing the CRISPRi lentiviral vector.

This work was supported by the foundation de France, France Parkinson, Edmond & Lily Safra foundation and the Institut Hospitalo-Universitaire de Neurosciences Translationnelles de Paris, A-ICM, Investissements d’Avenir ANR-10-IAIHU-06. Jana Heneine received funding from the Ministère de la Recherche et de l’Enseignement Supérieur and from the Fondation de la recherche Médicale (FRM, 4th year PhD program). The authors declare no competing financial interests.

## Authors contribution

JH performed most of the experiments. CCS and CPG helped with cell culture. CCS helped with RT-qPCR experiments, designed the lentiviral constructions for the knock-down experiments, transfected and FACS-purified the homogeneous cell pools containing the viral constructions. CZ et NA helped with the immunofluorescence experiments to investigate mitochondrial turnover, and BG helped for the image analyses from immunofluorescence experiments. JG, FXL and TG contributed to the bioinformatics and statistical analyses. OC helped with the conception of the study. JH, PR and HC conceived the study, designed the experiments, analyzed the data and wrote the manuscript. All authors read, checked and suggested modifications to the manuscript.

