## supplementary figure legends for "The UPR^ER^ governs the cell-specific response of human dopaminergic neurons to mitochondrial stress"

**Supplementary Figure 1. (a)** Detection of incorporated EdU (green) in DA neurons after incubation with EdU for 4 h or 24 h in the presence or absence of ddC. The right panels are zoomed areas from the left panels, indicated by dotted squares. **(b)** TOMM20 (red) expression assessed by immunofluorescence and EdU (green) detection in control conditions or following mitochondrial stress for 8 h. **(a, b)** Nuclei were stained using DAPI (blue). **(c)** Percentage of EdU-positive puncta co-localized with TOMM20 per neuron in control conditions and upon a 2 h-, 4 h- or 8 h-stress. Data were obtained from 3 independent differentiation experiments. Each dot represents the percentage of EdU puncta TOMM20 positive for one experiment of differentiation. The bar represents the mean of the 3 values, and the error bars show standard error of the mean.

**Supplementary Figure 2. (a)** Number of ATAC-seq peaks altered upon a 8 h long stress and defining chromatin regions with decreased or increased accessibility, depending on their genomic loci. **(b)** Gene ontology analysis (Biological Process 2023, Cistrome DB) performed on ATAC-seq peaks linked to promoters and displaying increased accessibility upon 8 h of stress compared to control conditions. **(c)** Gene ontology analysis (Cellular Component 2023, Cistrome DB) performed on ATAC-seq peaks linked to promoters and displaying decreased accessibility upon 8 h of stress compared to control conditions. **(d)** Gene ontology analysis (Biological Process 2023, Cistrome DB) performed on all ATAC-seq peaks displaying altered accessibility (increased in green, decreased in blue) upon 8 h of stress compared to control conditions.

**Supplementary Figure 3.** Phospho-Serine 65 ubiquitin (red), TOMM20 (green) and MAP2 (grey) expression in DA neurons treated with DMSO (Control) or after exposition to mitochondrial toxins for 4 h and 6 h (Stress), in the presence or absence of GSK2606414, observed by immunofluorescence. Nuclei were stained using DAPI (blue). For each time point, the right panels are zoomed areas from the left panels, indicated by dotted squares.

**Supplementary Figure 4. (a)** *Lnc-SLC6A15-5* expression, assessed by RT-qPCR, in DA neurons transduced by lentiviral vectors carrying *dCAS9-KRAB* and sgRNAs either targeting *lnc-SLC6A15-5* (KD *lnc-SLC6A15-5*) or a non-human sequence (NEG), in control conditions or following 8 h of mitochondrial stress (Two-way ANOVA with Tukey's multiple comparison test). RNA expression was normalized relatively to *TBP* mRNA expression. Data from 3 independent differentiation experiments, represented by 3 dots, were used. The bar represents the mean of the 3 values, and the error bars show standard error of the mean . **(b)** TH (green) and DAT (red) expression, assessed by immunofluorescence on DA neurons expressing normal (NEG) or reduced levels (KD) of *lnc-SLC6A15-5* in control conditions or upon 8 h mitochondrial stress . **(c)** Graphs show quantification of the percentage of $TH^+$ cells and $TH^+$ $DAT^+$ cells in both control and stress (8 h) condition (Two-way ANOVA with Tukey's multiple comparison test; no significant p-values found). **(d)** Phospho-Serine 65 ubiquitin (red), TOMM20 (green) expression in DA neurons expressing normal (NEG) or reduced levels (KD) of lnc-SLC6A15-5 and treated with DMSO (Control) or after 8 h long exposition to mitochondrial toxins (Stress), observed by immunofluorescence. **(e)** The graph represents the percentage of phospho-Serine65 ubiquitin-positive neurons in DA neurons expressing normal (NEG) or reduced levels (KD) of *lnc-SLC6A15-5*, in control conditions and following 8 h of mitochondrial stress (Two-way ANOVA with Tukey's multiple comparison test). **(f)** TOMM20 (red) and MAP2 (grey) expression assessed by immunofluorescence and EdU (green) detection in DA neurons expressing normal (NEG) or reduced levels (KD) of *lnc-SLC6A15-5* and treated with DMSO (Control) or after a 8 h long exposition to mitochondrial toxins (Stress). **(g)** Percentage of EdU-positive puncta in DA neurons expressing normal (NEG) or reduced levels (KD) of *lnc-SLC6A15-5*, in control conditions and following 8 h of mitochondrial stress (Two-way ANOVA with Tukey's multiple comparison test). **(b, d, f)** Nuclei were stained using DAPI (blue). The right panels are zoomed areas from the left panels, indicated by dotted squares. **(c, e, g)** Data were obtained from 3 independent differentiation experiments, represented by 3 dots. The bar represents the mean of the 3 values, and the error bars show standard error of the mean. *p-value ≤ 0,05; **p-value ≤ 0,01 ns p-value

**Supplementary Figure 5. (a)** Detection of OPP (grey) in DA neurons expressing normal (NEG) or reduced levels (KD) of *lnc-SLC6A15-5*, in control conditions or following 8 h of mitochondrial stress. Nuclei were stained using DAPI (blue). The graph displays the mean intensity of the OPP signal per $TH^+$ neurons, in control conditions or following 8 h of mitochondrial stress (Two-way ANOVA with Tukey's multiple comparison test). Data were obtained from 2 independent differentiation experiments. Each dot represents the mean intensity of the OPP signal in DA neurons for one experiment of differentiation. The bar represents the mean of the 2 values, and the error bars show standard error of the mean. **(b)** EIF2α, phosphorylated EIF2α (P-EIF2α) and Vinculin expression, assessed by Western Blot, in DA neurons expressing normal (NEG) or reduced levels (KD) of *lnc-SLC6A15-5*, under 4 different experimental settings. Quantification of the P-EIF2α/EIF2α ratio in DA neurons expressing normal (NEG) or reduced levels (KD) of *lnc-SLC6A15-5,* in control conditions and upon mitochondrial stress (8 h) from 2 independent differentiation experiments represented by 2 dots. The bar represents the mean of the 2 values, and the error bars show standard error of the mean. **(c)** Control (blue) and stress (green) conditions were performed either during 8 h (plain bars) or during 7h30 followed by 30 min recovery (hatched bars). *DDIT4, SYNCRIP, MTOR, RPS6KB1, RPS6, EIF4EBP1 and EIF4EBP2* mRNA expression, assessed by RT-qPCR, in DA neurons expressing normal (NEG) or reduced levels (KD) of *lnc-SLC6A15-5*, in the 4 experimental conditions (Type II Wald Chi-square tests ANOVA function with Tukey's multiple comparisons test). mRNA expression was normalized relatively to *TBP* mRNA expression. Data from 3 independent differentiation experiments, represented by 3 dots, were used. The bar represents the mean of the 3 values, and the error bars show standard error of the mean. *p-value ≤ 0,05; **p-value ≤ 0,01; ***p-value ≤ 0,001; **** p-value ≤ 0,0001.
