## Supplementary figures and images for "The UPR^ER^ governs the cell-specific response of human dopaminergic neurons to mitochondrial stress"

### Supplementary figure 1

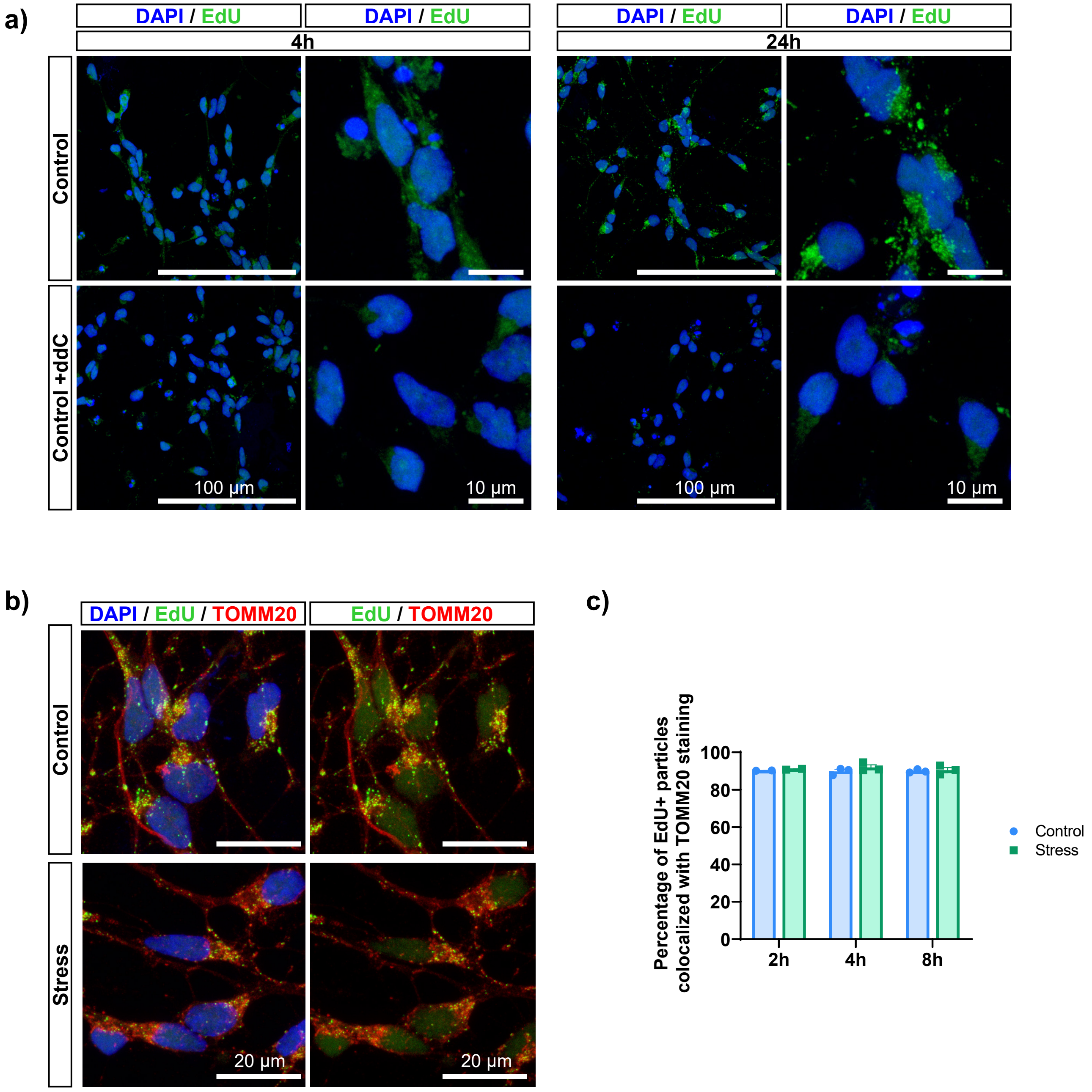

### Supplementary figure 2

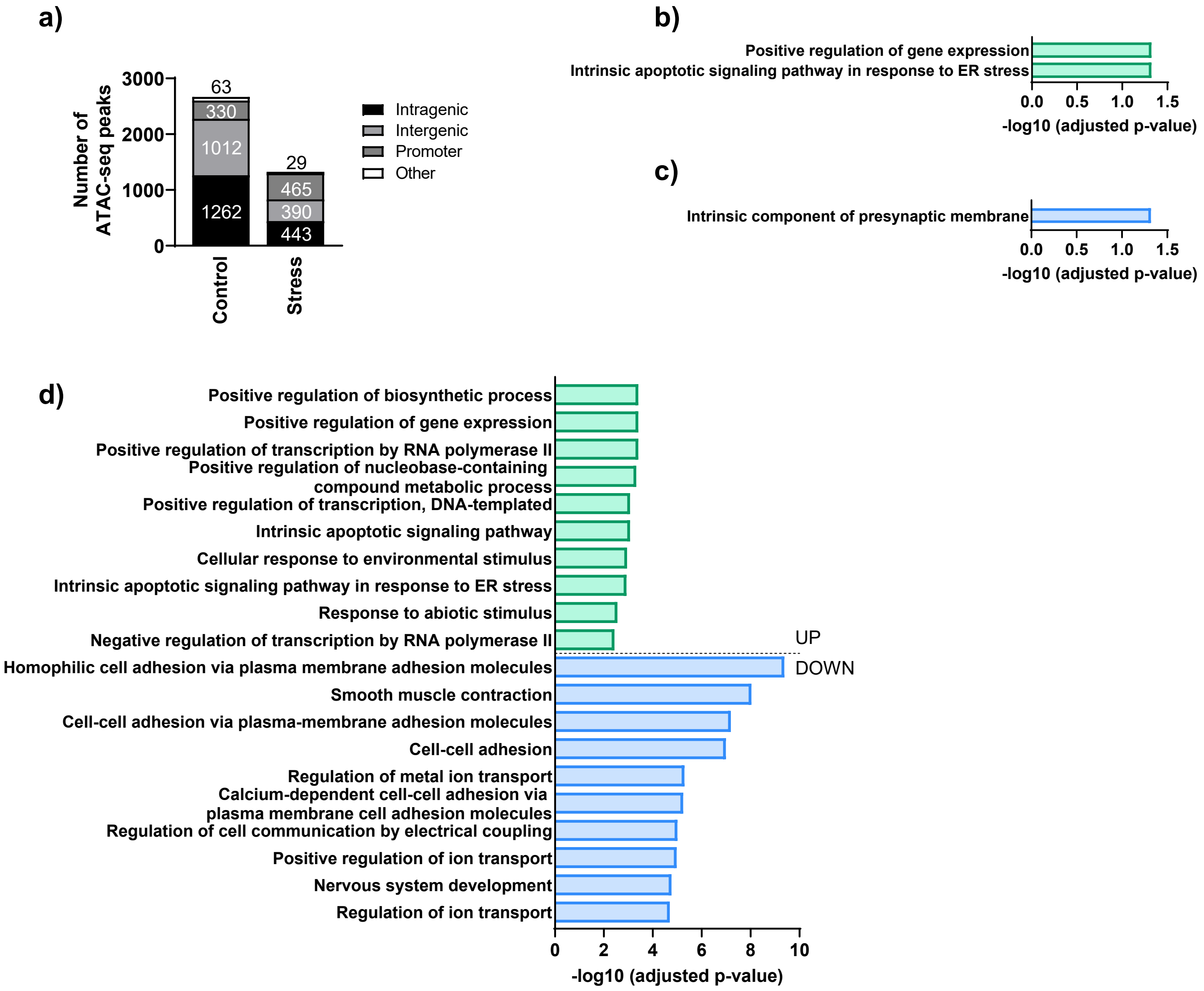

### Supplementary figure 3

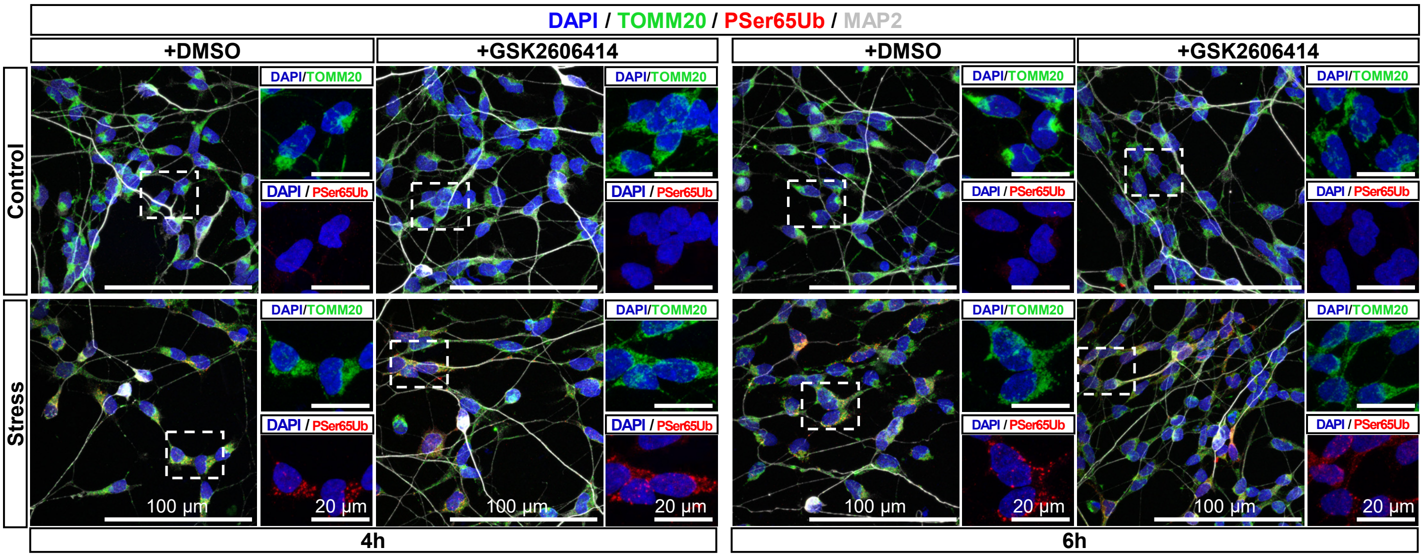

### Supplementary figure 4

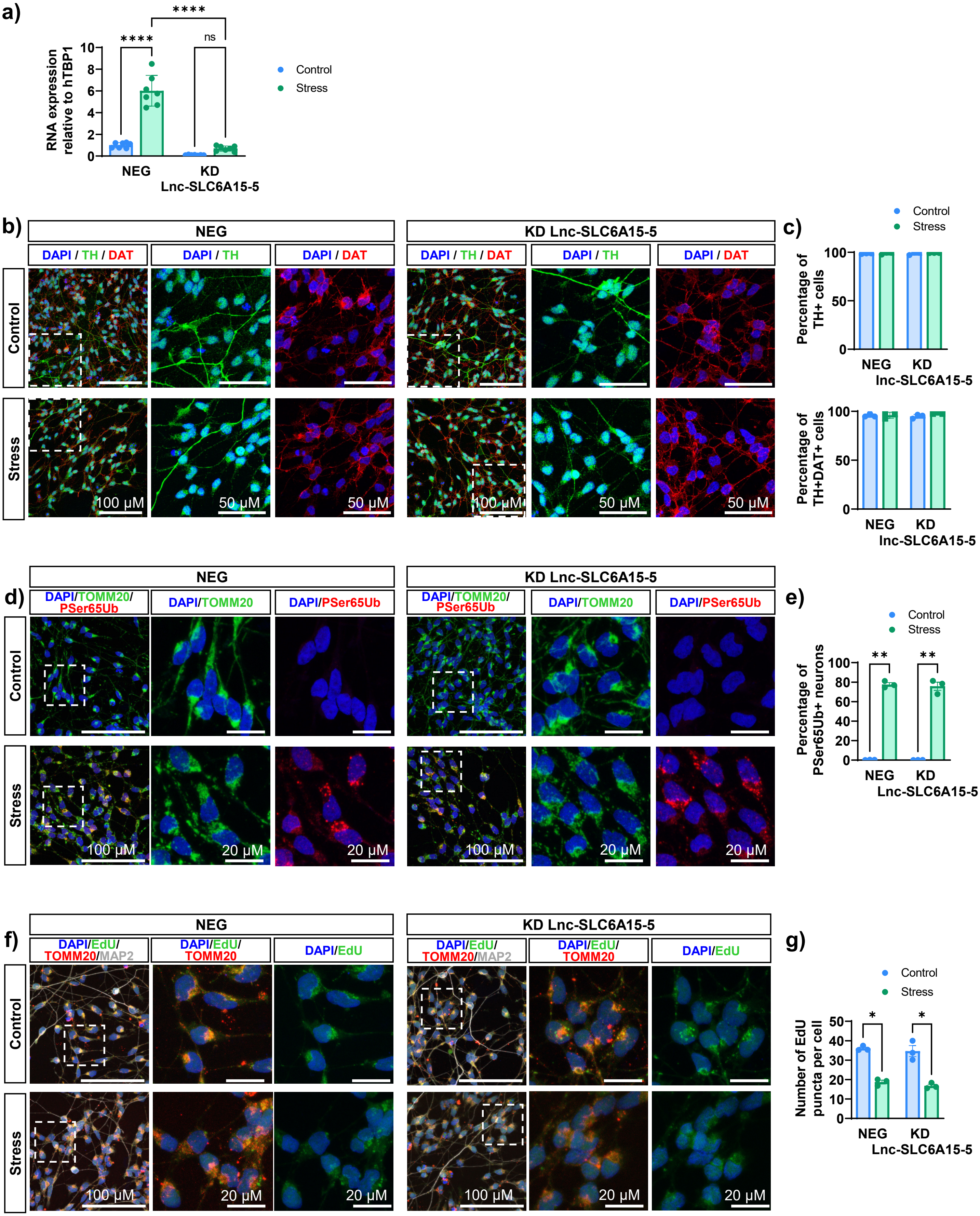

### Supplementary figure 5

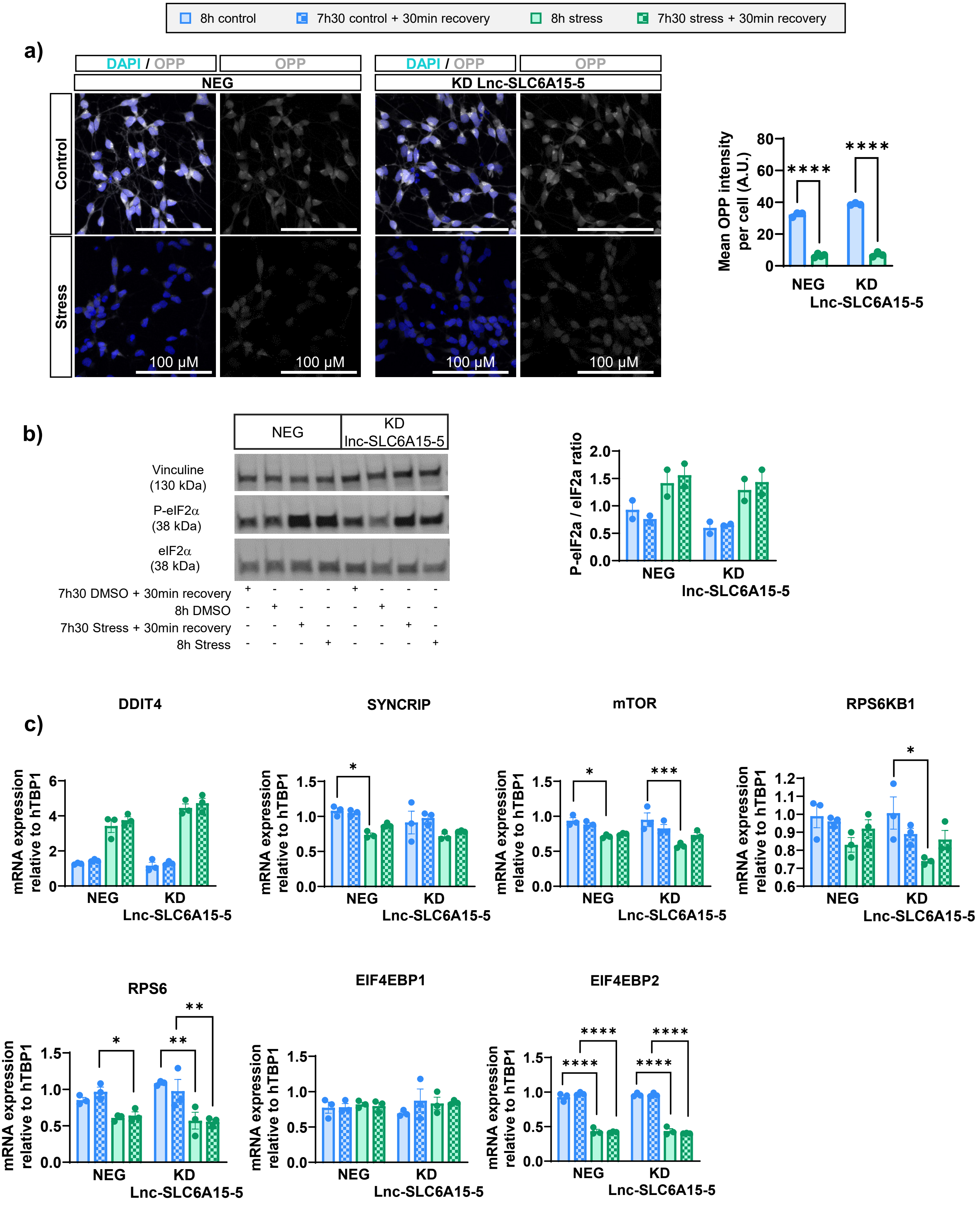
