## Supplementary Table S1 for "The UPR^ER^ governs the cell-specific response of human dopaminergic neurons to mitochondrial stress"

| Primary antibodies for Immunostaining | | | | |
|---|---|---|---|---|
| **Antigen** | **Supplier** | **Reference** | **Host** | **Dilution** |
| CASPASE 3 (Cleaved) | Cell Signaling Technology | 9661 | Rabbit | 1/500 |
| DAT | Millipore | mab369 | Rat | 1/400 |
| TH | Millipore | ab152 | Rabbit | 1/400 |
| Phospho Ser65 Ubiquitin | Cell Signaling Technology | N-70973S | Rabbit | 1/500 |
| TOMM20 | Abcam | ab56783 | Mouse | 1/1000 |
| MAP2 | Abcam | ab5392 | Chicken | 1/2500 |

| Primary antibodies for Western blotting | | | | |
|---|---|---|---|---|
| **Antigen** | **Supplier** | **Reference** | **Host** | **Dilution** |
| ATF4 (D4B8) | Cell Signaling | 11815S | Rabbit | 1/1000 |
| Phospho-EIF2α (Ser51) | Cell Signaling | 3398S | Rabbit | 1/1000 |
| EIF2α (D7D3) | Cell Signaling | 5324S | Rabbit | 1/1000 |
| Vinculin | Invitrogen | MA5-11690 | Mouse | 1/1000 |

| Secondary antibodies | | | | |
|---|---|---|---|---|
| **Antigen** | **Supplier** | **Reference** | **Host** | **Dilution** |
| Alexa Fluor Anti-Rabbit 555 | Thermo Fisher Scientific | A21428 | Goat | 1/1000 |
| Alexa Fluor Anti-Rat 555 | Thermo Fisher Scientific | A21434 | Goat | 1/1000 |
| Alexa Fluor Anti-Rabbit 488 | Thermo Fisher Scientific | A11070 | Goat | 1/1000 |
| Alexa Fluor Anti-Rabbit Cy3 | Thermo Fisher Scientific | A10520 | Goat | 1/1000 |
| Alexa Fluor Anti-Mouse 488 | Thermo Fisher Scientific | A11029 | Goat | 1/1000 |
| Alexa Fluor Anti-Mouse 555 | Thermo Fisher Scientific | A21422 | Goat | 1/1000 |
| Alexa Fluor Anti-Chicken 647 | Thermo Fisher Scientific | A21449 | Goat | 1/1000 |
| Peroxidase AffiniPure Anti-Rabbit | Jackson | 115-035-144 | Goat | 1/10000 |
| Peroxidase AffiniPure Anti-Mouse | Jackson | 115-035-003 | Goat | 1/20000 |
