## Supplementary Table S2 for "The UPR^ER^ governs the cell-specific response of human dopaminergic neurons to mitochondrial stress"

| Gene | Forward Primer | Reverse Primer |
|---|---|---|
| TBP | TGCACAGGAGCCAAGAGTGAA | CACATCACAGCTCCCCACCA |
| PERK | AGAGAGAGGAGCGTGTGTCT | TCCTGGTCCATTGCAGTCAC |
| EIF2A | GTTGCAACAGCTTATAGACCCC | GACAGTGTTTCGTGGTGTGC |
| ATF4 | TCCTCGATTCCAGCAAAGCA | CCAATCTGTCCCGGAGAAGG |
| ATF3 | CCTTTCATCTTCTTCAGGGGCT | AGGAAGAGCTGAGGTTTGCC |
| DDIT3 | CATGTTAAAGATGAGCGGGTGG | TGGATCAGTCTGGAAAAGCACA |
| CHAC1 | CGCTGTGGATTTTCGGGTAC | TTGCTTACCTGCTCCCCTTG |
| TRIB3 | GTGTCGCTTTGTCTTCGCTG | CTGCCTTGCCCGAGTATGAG |
| NRF2 | GAGCAAGTTTGGGAGGAGCT | GGTTGGGGTCTTCTGTGGAG |
| TRAF2 | CATACCCGCCATCTTCTCCC | TCATTCGGGCCCTTCATCAC |
| JNK | ATGAAGCTCTCCAACACCCG | GCCATTGATCACTGCTGCAC |
| ATF6 | TTCAGTCTCGTCTCCTCGGT | ATCTTCCTTCAGTGGCTCCG |
| HSPA5 | GACAAGAAGGAGGACGTGGG | GCATCGCCAATCAGACGTTC |
| EDEM1 | ATATGGTGCCCTCCCTGAGA | AGAAGCTCTCCATCCGGTCT |
| HERPUD1 | TTCCATTTAGACCGAGGCCG | GAGGTGGTTGGGGTCTTCAG |
| XBP1 | CGGAAGCCAAGGGGAATGAAG T | TGCAGAGGTGCACGTAGTCTGAG T |
| XBP1 s | CGGAAGCCAAGGGGAATGAAG T | ATACCGCCAGAATCCATGGGGAG A |
| IRE1 | CTGGAGCCTAGAGAAGCAGC | GGGACAGTGATGTTCTCCCG |
| DNAJC3 | ACCTGACAATGTGAATGCCCT | TGGAAGTTATCTGGGTGCCAC |
| ATF5 | GGATGGCTCGTAGACTATGGG | CGCTCAGTCATCCAGTCAGA |
| CLPP | ATGACATCTACTCGCGGCTG | TTGCAGGAAGAGGAGCTGTG |
| LONP1 | TGATCAACGTGTCGGGCTAC | CTTGGCCTTGCTCTCATCCA |
| YME1L1 | GGAGCCACAAACTTCCCAGA | AAGCCAACAGTACCTCGAGC |
| SIRT3 | CGTTGTGAAGCCCGACATTG | AAGTCCCGGTTGATGAGCAG |
| FOXO3 | TGGTTTGAACGTGGGGAACT | GCTGGGTTAGGAAAATGGCG |
| NRF1 | TCAGCAAACGCAAACACAGG | GTGACCGTGGTTGGCAATTC |
| lnc-SLC6A15-5 | GCAATGCTAGGCTCCTGACA | TATCCTCCCCGGGTTACTCG |
| VLDLR-AS1 | TACTTGCAGTTTCCAGGGGC | CACGTACGGCTTCTTTCTTGC |
| VPS11-DT | TGCCGGATGTGACTGTAACT | CCCTCATCTTGATCTCCCGG |
| lnc-FKRP | GAGTTTCTCTGGGTGGGACG | CAGCCCTCAGTGTCCAAGAC |
| SNHG1 | CGTTGGAACCGAAGAGAGCT | CTGTAACGCTGGCTTTGCAT |
| SNHG5 | CACAGTGGAGCAGCTCTGAA | GGCTACTCGTCCACACTCAG |
| TMEM161B-DT | TCTAAAGCAACTTCCGTGGGT | GCTGTTCGTCCCCTCACAAA |
| lnc-PSMC3IP-3 | ACATCCAGCTGCAACCTCTC | TGTATGCCACGTTGAGGGAC |
| MIR4697HG | GGAAAAGGCTCTGTCGTGGA | GAAGTGTGTGTGCAGGCTTG |
| FBXL19-AS1 | GGCTGTCCCCTCTATCCTCA | GGTGGAGAAGTGAGATGGGC |

| | | |
|---|---|---|
| NIPBL-DT | CGCTGGACAAGGCTGGAATA | TAGCGCACTGGTACACACAC |
| ZNF778-DT | TGTCTGAACATCACGCCGAA | TGGTCTTGGCTCCTACGGTA |
| lnc-MNAT1-2 | TCTTGCCTTTCCATGAGCGT | GGCAGAGGTGGATGGAGATG |
| lnc-TTC29 | GGAAAGGGGAGTGTTCACGT | ATGGCTCCTGATAACTGCGG |
| lnc-SLAIN1-11 | CGTGGCGCTAAGACTGAGAT | CCTTTCCCACCCCATTCGAA |
| SESN2 | TCTCCTCCTTCGTGTTTGGC | GGCTCTCTGACTTCTCCAGC |
| SLC3A2 | TGGTCCCAGTGGCGGATATA | CTACGGGGATGAGATTGGCC |
| SLC1A5 | CTCCTTGATCCTGGCTGTGG | GGGCAGCTCACTCTTCACTT |
| SLC7A5 | TTCTTCAACTGGCTCTGCGT | GAGACGGCGATCAGGAAGAG |
| SLC1A3 | ACATGAAGGAACAGGGGCAG | GGAGTGGCAAGACGATGACT |
| DDIT4 | GGTTCGCACACCCATTCAAG | CCAAAGGCTAGGCATGGTGA |
| SYNCRIP | TGGGAAACTGGAACGAGTGAA | TGATCTGGTGGCTTGGCAAA |
| mTOR | AGCTCTTCGGCCTGGTTAAC | CTTCTCCCTGTAGTCCCGGA |
| RPS6KB1 | ACTTCTGGCTCGAAAGGTGG | TGTTCGTGGGCTGCCAATAA |
| RPS6 | GCCCCAAAAGAGCTAGCAGA | GCAGGACACGTGGAGTAACA |
| EIF4EBP1 | CCTTCCAGTGATGAGCCCC | GTGTTCACGAAGAGGAGGGG |
| EIF4EBP2 | GGATCGTCGCAATTCTCCCA | GCCAAATCAGGTGCACACAA |
